# Deciphering the conformational dynamics of gephyrin-mediated collybistin activation

**DOI:** 10.1101/2022.05.30.493832

**Authors:** Nasir Imam, Susobhan Choudhury, Katherina Hemmen, Katrin G. Heinze, Hermann Schindelin

## Abstract

Efficient neuronal signaling depends on the proper assembly of the postsynaptic neurotransmitter machinery and at inhibitory GABAergic synapses is controlled by the scaffolding protein gephyrin and collybistin, a Dbl-family guanine nucleotide exchange factor and neuronal adaptor protein. Collybistin usually contains an N-terminal SH3 domain and exists in closed/inactive or open/active states. Here, we elucidate the molecular basis of the gephyrin-collybistin interaction with newly designed collybistin FRET sensors. Using fluorescence lifetime-based FRET measurements, we deduce the affinity of the gephyrin-collybistin complex, thereby confirming that the C-terminal dimer-forming E domain binds collybistin, an interaction, which does not require E domain dimerization. Simulations based on fluorescence lifetime and sensor distance distributions reveal a dynamic behavior of the SH3 domain already in the closed state of collybistin. Finally, our data provide strong evidence for a collybistin-gephyrin communication network, where, unexpectedly, switching of collybistin from closed/inactive to open/active states is efficiently triggered by gephyrin.

## Introduction

Inhibitory synaptic signaling in the mammalian brain heavily depends on the neurotransmitter γ-aminobutyric acid (GABA), which is recognized by postsynaptic GABA type-A (GABA_A_) receptors. Postsynaptic GABA_A_ receptor clustering in direct apposition to the presynaptic transmitter release sites ensures fast signal transduction, inducing postsynaptic membrane hyperpolarization and reduced excitability (Andersen et al., 1963; Buhl et al., 1994; Miles & Wong, 1984; Nusser et al., 1997). GABA_A_ receptor assembly and maintenance is coordinated by various postsynaptic neuronal factors. These core neuronal components include cell adhesion proteins of the neuroligin family, the scaffolding protein gephyrin and the guanine nucleotide exchange factor (GEF) collybistin (Luscher et al., 2011; Papadopoulos & Soykan, 2011).

Gephyrin serves as a prime scaffolding protein at inhibitory GABAergic and glycinergic postsynaptic specializations and is principally responsible for GABA_A_ and glycine receptor clustering (Betz, 1998; Choii & Ko, 2015; Fritschy et al., 2008; Tyagarajan & Fritschy, 2014). Gephyrin splice variants encompass two structured domains; a trimeric N-terminal G-domain, and a C-terminally located dimerizing E-domain, which are separated by a long unstructured linker (Choii & Ko, 2015; Kim et al., 2006; Pizzarelli et al., 2020; Sola et al., 2004). Based on the oligomeric states of the isolated gephyrin G and E domains, a hexagonal lattice model has been suggested to provide a framework for gephyrin-mediated molecular organization at inhibitory synapses (Crosby et al., 2019; Tyagarajan & Fritschy, 2014; Xiang et al., 2001). However, in full-length gephyrin only G-domain trimerization takes place, whereas E-domain dimerization is prevented, predominantly resulting in gephyrin trimers (Sander et al., 2013). Gephyrin loss is lethal and causes mice to die within the first post-natal day (Feng et al., 1998). Previous studies (Jedlicka et al., 2009; Papadopoulos et al., 2008; Papadopoulos et al., 2007) demonstrated that clustering of GlyRs and GABA_A_Rs in gephyrin-deficient mice is markedly reduced, underlining the essential role of gephyrin in receptor assembly at inhibitory postsynapses. Gephyrin mutations have also been associated with various brain disorders, including autism, schizophrenia, Alzheimer’s disease, and epilepsy (Agarwal et al., 2008; Dejanovic et al., 2014; Förstera et al., 2010; Kiss et al., 2016; Lionel et al., 2013).

Gephyrin recruitment from intracellular deposits to postsynaptic membranes mainly depends on the adaptor protein collybistin (CB; also referred to as ARHGEF9). As a Dbl-family member CB contains tandem Dbl-homology (DH) and pleckstrin-homology (PH) domains (Zheng, 2001). In addition to the DH domain catalyzing the GEF activity and the phosphoinositide-binding PH domain, the majority of CB mRNAs encode an additional N-terminal src-homology 3 (SH3) domain (Harvey, 2004). In rat, CB genes are expressed in the splice variants CB1, CB2 and CB3, which differ in their N and C termini and the presence or absence of the SH3 domain (Harvey, 2004; Kins et al., 2000). Structural and biochemical studies suggest that the most abundantly expressed, full-length, SH3-domain-containing CB isoform 2 (CB2-SH3^+^), adopts a closed conformation wherein the N-terminal SH3 domain interacts intra-molecularly with residues in the DH and PH domains, a conformation which does not interact with Cdc42 (Soykan et al., 2014). In contrast, the SH3 domain lacking CB2 splice variant (CB2-SH3^-^) constitutively regenerates the GTP-bound state of Cdc42 (Reddy-Alla et al., 2010; Tyagarajan et al., 2011; Xiang et al., 2006). Previous studies indicated that CB-mediated gephyrin recruitment and clustering at the plasma membrane depend on binding of its PH domain to phosphatidylinositol 3-phosphate [PI(3)P], whereas the GEF activity of its DH domain is dispensable (Kalscheuer et al., 2009; Reddy-Alla et al., 2010). Mutations causing a disruption of CB inter-domain association lead to an open/active (with respect to receptor anchoring) CB conformation with enhanced phosphatidylinositol affinity (Soykan et al., 2014). *In vivo* experiments with CB-deficient mice indicated a reduction in synaptic gephyrin and GABA_A_ receptor γ2-subunit clustering, decreased GABAergic synaptic transmission and impaired spatial learning (Papadopoulos et al., 2008). Surprisingly, CB-deficient mice do not exhibit deficits in glycinergic synaptic transmission, suggesting that CB is dispensable for gephyrin-mediated GlyR clustering at glycinergic synapses, but is required for the clustering of certain GABA_A_ receptors (Körber et al., 2012; Saiepour et al., 2010), further stressing the vital role of CB in the initial assembly and maintenance of gephyrin-GABA_A_ receptor clusters. However, owing to technical limitations, clear molecular insights into the association of CB and gephyrin and the overall clustering process have been lacking. Although the CB-gephyrin interaction has already been reported in previous studies (Kins et al., 2000; Tyagarajan et al., 2011), no quantification of the interaction strength has been reported and the reciprocal regulation of the activities of both proteins remain poorly understood.

The present study aims at elucidating the molecular basis of CB autoinhibition, its binding to gephyrin and whether this interaction leads to CB activation. We constructed novel intramolecular CB FRET sensors to understand the conformational dynamics of CB by making use of picosecond-scale time-resolved fluorescence lifetime measurements. These studies confirmed that CB in isolation remains in a closed conformation while gephyrin binding takes place via its C-terminal E domain leading to an open conformation of CB. We quantified the interaction strength of gephyrin to wild type and constitutively active, open state mutant CB FRET sensors. Based on the inter-fluorophore distance distributions from the FRET sensors in the absence and presence of gephyrin, we modeled the overall three-dimensional conformational space accessible to CB with respect to the SH3 domain. Our data combined with simulation studies provide clear molecular evidence of gephyrin-mediated CB opening, thus suggesting a synergy between concurrent gephyrin scaffold assembly and CB targeting to the plasma membrane.

## Results

### Sensor engineering and characterization

To generate suitable FRET sensors we incorporated an *Aequorea victoria* derived cyan variant (CFP) of the green fluorescent protein (Heim et al., 1994) and the biarsenical dye, fluorescein arsenical hairpin binder-ethanedithiol (FlAsH) (Griffin et al., 1998), as a suitable FRET pair (Hoffmann et al., 2005) into CB (Fig. 1A-B). The non-fluorescent molecule FlAsH forms a fluorescent complex with any protein to which a short tetracysteine-motif (tCM) is genetically fused possessing the highest specificity for the amino acid sequence CCPGCC (Fig. 1C, Table S1) in the binding motif (Adams et al., 2002). We optimized the position for CFP attachment and the tCM insertion site so that the intramolecular CB-FRET sensor can be used to study ligand-induced conformational dynamics. Initial screening for suitable intramolecular CB-FRET sensors was done by attaching the CFP moiety at various C-terminal positions by utilizing different truncated forms of CB, while the tCM insertion site was kept constant at the N-terminus of CB. Insertion of tCM after the first amino acid residue and CFP after residue 456 of the CB2-SH3^+^ splice variant (Fig. 1B, Table S1), denoted as F1, yielded the best working wild-type CB FRET sensor in terms of FRET efficiency (E_FRET_) along with sensor purity, while still maintaining proper folding (Fig. S1, A-B). To better understand CB conformational dynamics, we generated three additional CB-FRET sensors (Fig. 1B) where we inserted the tCM for the FlAsH labeling after residues 28 (F28), 73 (F73) and 99 (F99) of CB while keeping the CFP position fixed after residue 456. Additionally, we also constructed two open state mutant CB sensors (Fig. 1A-B). In the single mutant sensor, (smCB), Glu262 was replaced with alanine, whereas in the double mutant sensor (dmCB), an additional Trp24Ala variant was engineered while keeping the FlAsH and CFP at the same positions as in the F1 construct.

**Figure 1.**
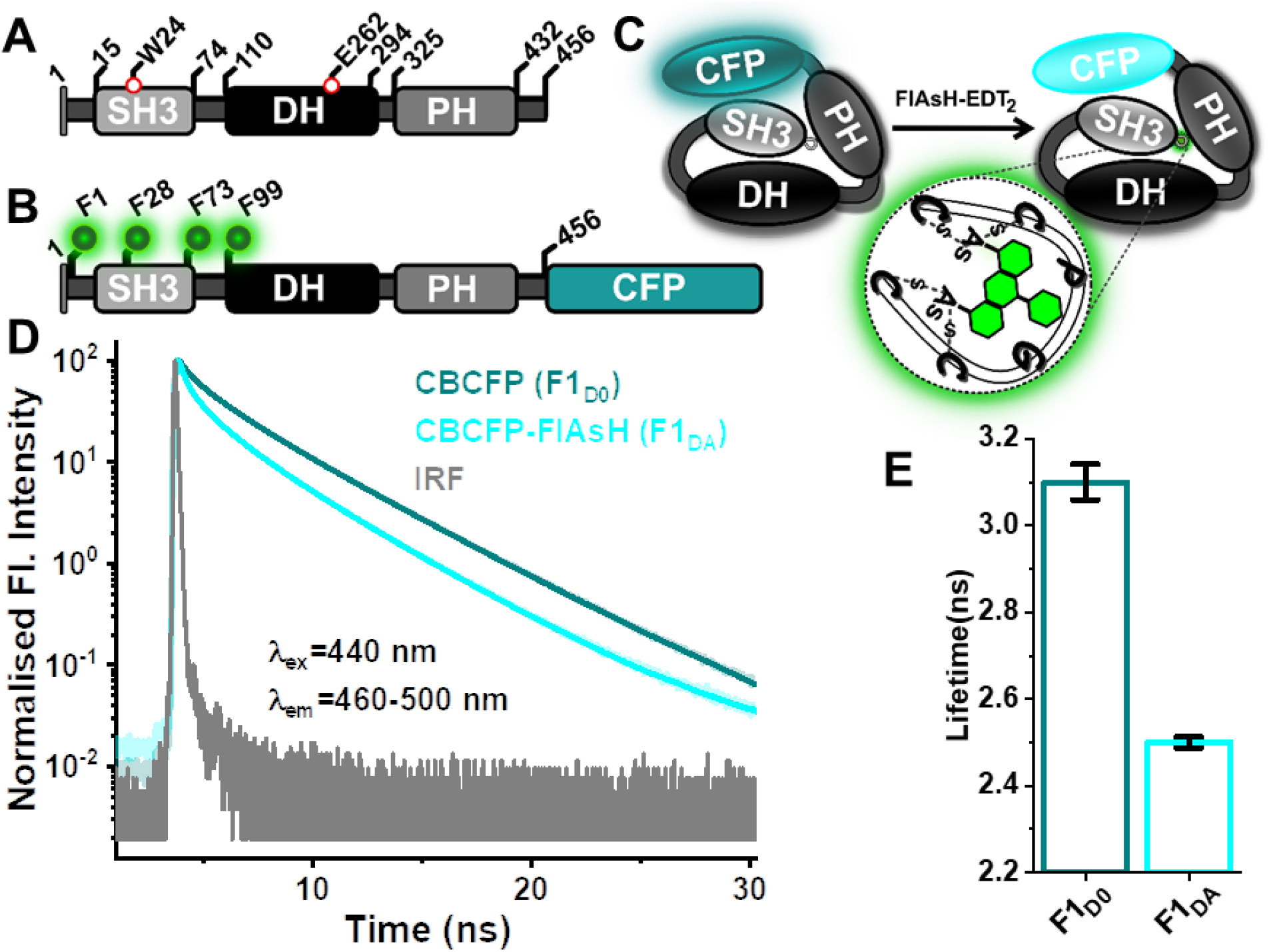
CB FRET sensor design and characterization. **(A)** CB domain architecture with the SH3, DH and PH domains in light grey, black and dark grey, respectively. Amino acid positions Trp24 (W24) and Glu262 (E262) are highlighted. **(B)** Domain architecture of the ensemble of CB FRET sensors constructed in this study, highlighting the position of the tetra-Cysteine motif (tCM) used for labeling with the fluorescein arsenical hairpin binder-ethanedithiol (FlAsH-EDT2) (green spheres) and C-terminal attachment site of CFP (teal). Individual sensors contained a solitary tCM inserted after residue 1 (F1), 28 (F28), 73 (F73) and 99 (F99), whereas the CFP position (after residue 456) was kept constant. F1_D0_, F28_D0_, F73_D0_ and F99_D0_ represent the individual FRET sensors in the absence of FlAsH and F1_DA_, F28_DA_, F73_DA_ and F99_DA_ after FlAsH-labeling. In the single mutant FRET sensor (F1_smDA_) E262 was replaced with Ala (E262A) and in the double mutant FRET sensor (F1_dmDA_) W24 and E262 were replaced with Ala (W24A/E262A). **(C)** Cartoon representing the CB FRET sensor (F1_D0_) in the closed conformation highlighting its labeling with FlAsH-EDT_2_ reagent, resulting in F1_DA_. FlAsH-EDT_2_ (non-fluorescent) turns fluorescent (green) after forming covalent bonds with the cysteine residues present in tCM (green circle). **(D)** Time-resolved fluorescence intensities of CFP of the CB FRET sensor (F1_D0_; teal) and the FlAsH-labeled CB FRET sensor (F1_DA_; cyan). Instrument response function (IRF) is shown in grey. F1_D0_ and F1_DA_ were excited (λ_ex_) at 440 nm and emission (λ_em_) data were collected between 460-500 nm. Data were scaled to a maximum of 100 for easier comparison. **(E)** Species-weighted ⟨τ⟩ of CFP in F1_DA_ (2.5±0.02 ns) is reduced compared to F1_D0_ (3.1±0.04 ns), corresponding to a FRET efficiency of 0.19 (eq. 8). Data from three different batches of experiments are presented as mean values ± SD.

Previous crystallographic studies (Soykan et al., 2014) suggested that CB exists in a closed conformation in its inactive state which would allow optimum resonance energy transfer between CFP and FlAsH (Fig. 1C). For our initial studies we used the F1 construct which is expected to closely mimic wild type CB2-SH3^+^. Time-resolved fluorescence intensities of CFP in the absence (F1_D0_) and presence of FlAsH (F1_DA_) revealed a significant reduction in the average fluorescence lifetime (⟨τ⟩) from 3.1 ± 0.04 ns to 2.5 ± 0.02 ns (Fig. 1D-E, Table S3). The decrease in ⟨τ⟩ in F1_DA_ is attributed to FRET from the C-terminally attached CFP to the FlAsH moiety bound to tCM. F1_DA_ displayed a FRET efficiency (E_FRET_) of ∼19% (eq. 6) and a calculated Förster radius (R_0_) of 39 Å (eq. 6 and Fig. S1 C). These properties support the use of F1_DA_ as FRET sensor to study ligand-induced CB conformational dynamics.

### Gephyrin, NL2 and SH3 mediate CB activation

Full-length CB is rather unstable and prone to degradation, while full-length gephyrin (GephFL) is susceptible to aggregation and degradation during purification (Sander et al., 2013; Soykan et al., 2014). After optimization, we recombinantly purified stable constructs of full-length CB and GephFL (see Methods for details). Using the CB FRET sensor as a novel tool, we sought to delineate the molecular basis of the CB and GephFL interaction. For interaction studies, we measured the ⟨τ⟩ of CFP in F1_D0_ and F1_DA_ in the presence of a 100-fold molar excess of GephFL (100 µM). A significant increase in average fluorescence lifetime to 2.73 ± 0.02 ns (Table S3) of the F1_DA_-GephFL complex compared to free F1_DA_ was observed (Fig. 2A). An E_FRET_ of ∼12% was calculated for the F1_DA_-GephFL complex (Table S3), while no significant change in ⟨τ⟩ of CFP was observed for F1_D0_ in the presence of GephFL (Fig. S3A), indicating that GephFL binding does not alter the fluorescent properties of CFP. The decrease in E_FRET_ upon GephFL binding reflects changes in the CB conformation, leading to an increased distance between the SH3 and PH domains.

**Figure 2.**
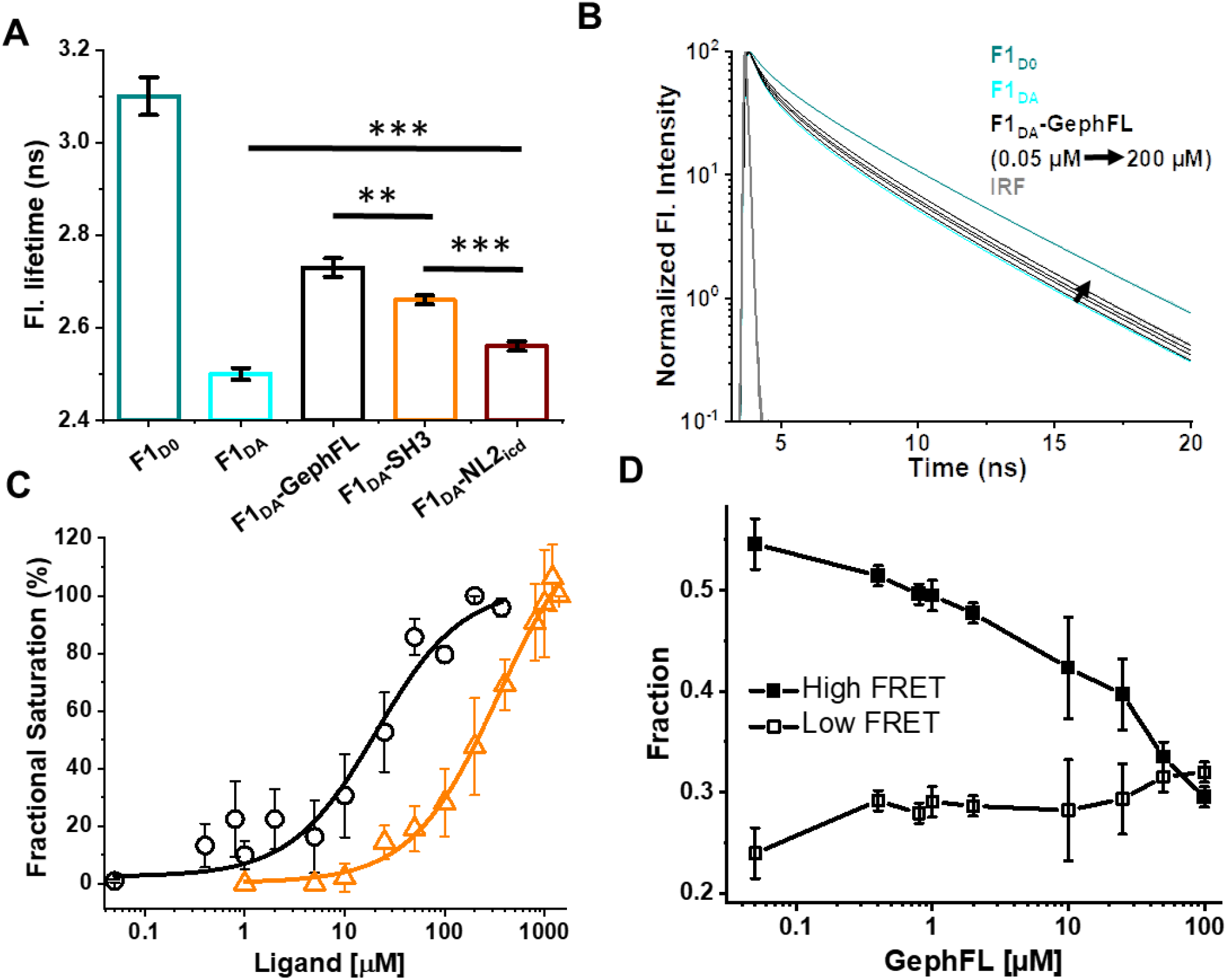
Full-length gephyrin-mediated CB activation. **(A)** Average fluorescence lifetime of CFP in F1_D0_ (teal), F1_DA_ alone (cyan) and in the presence of GephFL (black), SH3 domain (orange) and NL2_icd_ (dark red). *** P< 0.001, ** P< 0.01 and P< 0.05. **(B)** Fluorescence lifetime of CFP in F1_DA_ and F1_DA_-GephFL complexes with increasing concentrations of full-length gephyrin (GephFL, black). Data were scaled to a maximum of 100 for easier comparison. The decay histograms are fitted in two ways: i) With a multi-exponential fitting model (eq. 1) to determine K_D_ and ii) with two FRET species (R(i)) with a Gaussian distance distribution (half-widths 5 Å) and a single NoFRET species (see Methods). **(C)** GephFL binding affinity was determined based on the average fluorescence lifetime converted into the fractional saturation using eq. 4. The data were fitted with eq. 5. For the F1_DA_-GephFL complex (black) a dissociation constant (K_D_) of 20.2 ± 8.3 µM and for the F1_DA_-SH3 complex (orange) of 339 ± 44 µM (data ± SD) were obtained. Data from three different batches of experiments are presented as mean values ± SD. **(D)** Plot of the contribution of the two FRET species (high and low FRET states) against the concentration of GephFL obtained when analyzing the time-resolved fluorescence intensities with the Gaussian distribution model (eq. 9). Curves showing the fraction of F1_DA_ molecules in the closed/high FRET state (filled squares, black) with a FRET pair distance (R1) of 25 ± 1.1 Å and their gradual transition into the open/low FRET state (open squares) exhibiting a FRET pair distance (R2) of 46 ± 1.5 Å upon addition of GephFL. The remaining xNoFRET population also gradually increases upon addition of GephFL (Fig. S2C).

Previous studies hypothesized that the intracellular cytosolic domain of NL2 (NL2_icd_) binds to the SH3 domain of CB, resulting in an open/active state capable of interacting with plasma membrane phosphoinositides (Poulopoulos et al., 2009; Schäfer et al., 2020; Soykan et al., 2014). In the presence of 100 µM NL2_icd_ the ⟨τ⟩ of CFP in the F1_DA_-NL2_icd_ complex was 2.6 ± 0.02 ns (Fig. 2A) with an E_FRET_ of ∼16% (Table S3), a slightly higher E_FRET_ than in the F1_DA_-GephFL complex. To investigate the role of the SH3 domain during CB opening we incubated a 100-fold stoichiometric excess (100 µM) of recombinantly purified SH3 domain with F1_DA_. ⟨τ⟩ and E_FRET_ for the SH3-F1_DA_ complex were measured as 2.66 ± 0.02 ns and ∼14%, respectively (Fig. 2A, Table S3), indicating that the free SH3 domain can displace the covalently attached SH3 domain during CB opening.

Encouraged by these initial results, we performed titration experiments to quantify the binding affinity between CB and GephFL as shown in Fig. 2B. With increasing (0.05 to 200 µM) GephFL concentrations, a concomitant increase in F1_DA_ ⟨τ⟩ was observed, reaching saturation at a GephFL:F1_DA_ molar ratio of 100:1 (Fig. 2B-C). In contrast, when titrating with the free SH3 domain, saturation was only obtained at a SH3:F1_DA_ 1200:1 molar ratio (Fig. 2C and Fig. S3A). By plotting the fractional saturation determined from the corresponding ⟨τ⟩ (eq. 4) against the GephFL and SH3 concentrations, the dissociation constants (K_D_) of the F1_DA_-GephFL and F1_DA_-SH3 complexes were assessed as 20.2 ±8.3 µM and 339 ± 44 µM, respectively (eq. 5, Fig. 2 and Fig. S3). These results indicate that GephFL and CB interaction is moderately tight, while the SH3 domain has a low affinity towards CB. Previous microscale thermophoresis (MST) data (Soykan et al., 2014) with the free SH3 and the tandem DH-PH domains yielded a K_D_ of 273 ± 34 µM, similar to the value obtained here. The interaction strength between F1_DA_ and NL2_icd_ could not be quantified as no systematic increase in ⟨τ⟩ of CFP was observed when further incubated with excess (more than 100 µM) of NL2_icd_.

### Two state dynamics during CB activation

To understand the mode of CB activation, we analyzed the time-resolved fluorescence intensities of F1_DA_ and F1_DA_-GephFL complexes at various concentrations by Gaussian distance distribution models (eq. 9). From Fig. S2A it is evident that fitting with two FRET species with a Gaussian distance (R(i)) distribution and a NoFRET (x_NoFRET_) state is significantly better than assuming only a single FRET species, while three FRET species did not improve the fits further. The half-widths of the Gaussian distributions were kept fixed to 5 Å. The results suggested that the F1_DA_ molecules exist in two conformational states, a high-FRET and a low-FRET state (Fig. S2 B). The high FRET state exhibited an average inter-fluorophore distance (R1) of 25.5 ± 0.5 Å, while the low FRET state showed an average inter-fluorophore distance (R2) of 45.5 ± 0.9 Å (Table S3). Considering the size of CFP (diameter ∼20 Å), the distance of the high FRET state indicates that the FlAsH and CFP fluorophores are in very close proximity, in line with a compact/closed conformation, while in the low FRET state F1_DA_ adopts an open state. While gradual addition of GephFL did not induce any significant changes in the inter-fluorophore distances, it shifted the equilibrium towards the low FRET state (Fig. 2D). It must be noted that increasing GephFL concentrations led to a stronger population of the x_NoFRET_ state (Fig. S2 C), possibly indicating another state beyond the measurable FRET distance limit (>49 Å) for this FRET pair (Algar et al., 2019). The fluorescence lifetime-based FRET study along with distance distribution analysis of F1_DA_ provides concrete evidence of GephFL-mediated CB opening and its transition from the closed to an open state. Binding of the isolated SH3 domain also resulted in a concentration-dependent increase in the low-FRET F1_DA_ state and a simultaneous decline in the high-FRET F1_DA_ population (Fig. S3B-C), suggesting a displacement of the SH3 domain present in F1_DA_ by the isolated SH3 domain. Again, we observed that rising SH3 concentrations resulted in a x_NoFRET_ increase (Fig. S3D).

### The E domain of gephyrin mediates CB binding and activation

Next, we investigated which region of gephyrin mediates the interaction with CB by employing constructs containing only the G domain (GephG), the linker followed by the E domain (GephLE), and the isolated E domain (GephE) of gephyrin. The purified variants (GephG, GephLE and GephE) were incubated with F1_D0_ and F1_DA_, and fluorescence lifetime measurements were performed. Fig. 3A shows the ⟨τ⟩ changes in F1_DA_ when incubated with the domain variants. At comparable concentrations of 100 µM, GephLE showed the highest increase in ⟨τ⟩ with 2.67 ± ns, followed by GephE (2.59 ± 0.02 ns) (Fig. S4A, Table S3). In contrast, GephG (2.51 ± ns) did not display a fluorescence lifetime change, thus confirming previous studies reporting that the G domain is not involved in the interaction with CB (Tyagarajan & Fritschy, 2014; Tyagarajan et al., 2011). Note, the incubation of F1_D0_ with the domain variants, as observed before for GephFL, did not alter ⟨τ⟩ of CFP (Table S4). Taken together, these data suggest that GephE with a possible minor contribution from the linker mediates interaction and opening of CB.

**Figure 3.**
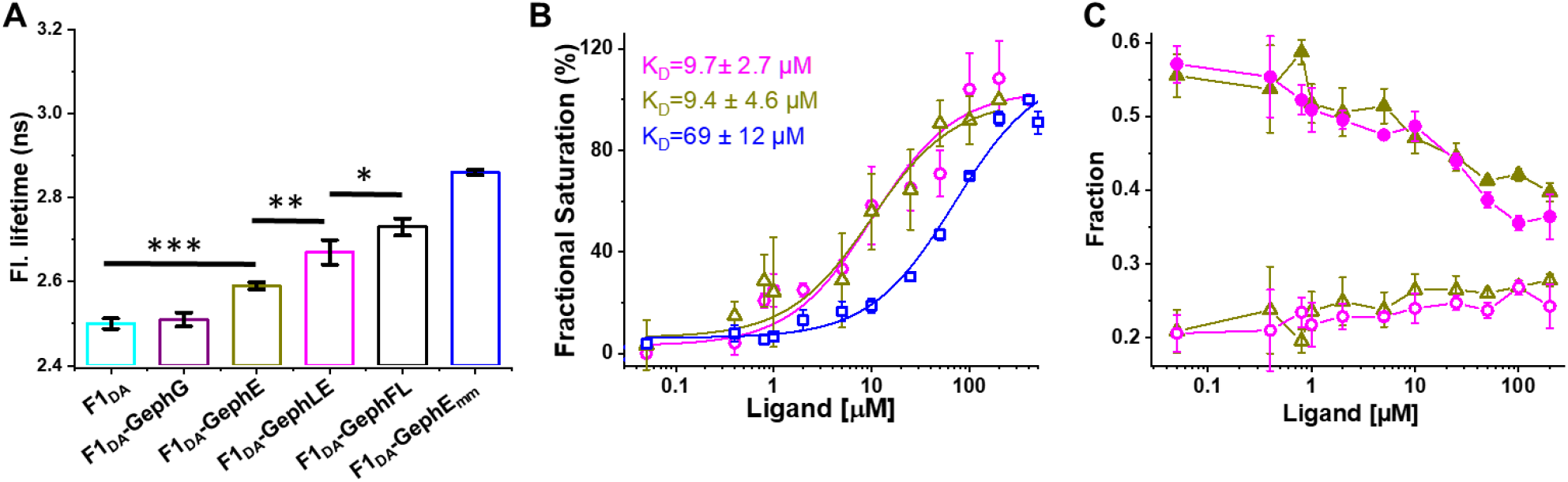
Gephyrin E domain mediates CB activation. **(A)** Bar graph showing the observed species-weighted average fluorescence lifetime of CFP in F1_DA_ (cyan) and in the presence of GephG (purple), GephE (dark yellow), GephLE (magenta) and the monomeric E domain variant (GephEmm, blue). *** P< 0.001, ** P< 0.01 and * P< 0.05. **(B)** Binding affinity curves for GephLE (magenta), GephE (yellow) and GephEmm (blue) determined based on the average fluorescence lifetime converted into fractional saturation using eq. 5. Curves are fitted with eq. 8 (see Methods) to determine the K_D_ for GephLE, GephE and GephEmm. Data from three different batches of experiments are presented as mean values ± SD. **(C)** Plot of the contribution of the two FRET species (high FRET state and low FRET state) against the concentrations of GephLE (triangle) and GephE (circle) obtained when analyzing the time-resolved fluorescence intensities with the Gaussian distribution model (eq. 9). The high FRET state (R1, filled circle and triangle) decreases, and the low FRET (R2, open circle and triangle) state increases with increasing concentrations of both ligands.

To further map the binding site, we individually titrated F1_DA_ with GephLE and GephE. F1_DA_ displayed a concentration-dependent increase in ⟨τ⟩ for both domain variants (Fig. S4 B-C). We quantified the binding strengths by plotting the fractional saturation based on the change in F1_DA_ ⟨τ⟩ (eq. 4) upon increasing GephLE and GephE concentrations, respectively. Both, GephLE and GephE displayed identical affinities with K_D_ values of 9.7 ± 2.7 µM and 9.4 ± 4.6 µM, respectively (Fig. 3B, eq. 5). Subsequently, we examined, how GephLE and GephE mediate the high-FRET to low-FRET transition in F1_DA_. In line with the identical affinities, GephLE and GephE displayed comparable concentration-dependent effects in transitioning from the high-FRET/closed state to the low-FRET/open F1_DA_ state (Fig. 3C, Fig. S4 D-E). Like GephFL, both GephLE and GephE exhibited an increase of x_NoFRET_ upon increasing the concentration of both ligands (Fig. S4 F-G). Hence, the fluorescence lifetime-based affinity interaction study with F1_DA_ demonstrated that the E domain solely mediates the interaction with CB.

### A monomeric E-domain induces a stronger CB conformational change

As GephE forms a dimer in its native state, we next checked whether GephE dimerization plays any role in its recognition by CB. To investigate this aspect we recombinantly purified a dimerization-deficient, monomeric mutant of GephE (GephE_mm_) described earlier (Saiyed et al., 2007) (Fig. S5 A). We then measured the ⟨τ⟩ change observed for F1_DA_ in the presence of GephE_mm_ and compared it to that of GephE (Fig. 3A). Surprisingly, at comparable concentrations, GephE_mm_ displayed a longer lifetime (2.85 ± 0.01 ns) compared to GephE (2.59 ± 0.02 ns), implying that GephE_mm_ possesses a higher potential for changing the conformation of CB (Table S3). To investigate the binding strength of GephE_mm_ we titrated F1_DA_ with increasing concentrations of GephE_mm_ and quantified the results (Fig. S5 B). Unexpectedly, GephE_mm_ displayed a significantly lower affinity with a K_D_ value of 69 ± 2.7 µM (Fig. 3B, eq. 5) compared to GephE (9.7 ± 2.7 µM). To better understand the conformational changes induced in F1_DA_ by GephE_mm_ we performed distance distribution fittings for both constructs as described for GephFL. GephE_mm_ was found to be more potent in turning the high FRET F1_DA_ molecules into a low FRET population compared to GephE/LE (Fig. 3C, Fig S5 C-D), however, the inter-fluorophore distance for the high FRET (R1) and low FRET (R2) molecules remained relatively unchanged for GephE (R1 = 23.6 ± 0.4 Å and R2 = 46.5 ± 1.8 Å) and GephE_mm_ (R1 = 24.5 ± 0.2 Å and R2 = 43.7 ± 0.3 Å) (Table S3). Hence, comparative fluorescence lifetime changes along with analyses of distance distribution results for GephE and GephE_mm_ clearly depict that GephE dimerization is not crucial for the CB-gephyrin interaction.

### Active state mutant sensor design and characterization

A previous study reported that residues Trp24 and Arg70, which are located in the SH3 domain, and Glu262 in the DH domain play crucial roles in modulating the equilibrium between the inactive and active conformations in full-length CB (Soykan et al., 2014). The W24A and E262A variants promote the formation of the open state, which, with respect to its role in inhibitory synapse formation, is considered to be the active state (Soykan et al., 2014). Small angle X-ray scattering (SAXS) and atomic force microscopy (AFM) data indicated a more extended conformation for the E262A single mutant, in contrast to the compact state of wild-type CB (Soykan et al., 2014). The W24A/E262A double mutant could not be analyzed by these biophysical techniques due to enhanced instability of the protein (Soykan et al., 2014). In our current study, however, we could carry out experiments with the single (F1_smDA_) as well as the double mutant (F1_dmDA_) CB FRET sensor.

The ⟨τ⟩ of CFP in F1_smD0_ (3.12 ± 0.02 ns) and F1_dmD0_ (3.13 ± 0.03 ns) in the absence of the FlAsH sensor were identical to that observed for F1_D0_ (3.10 ± 0.04 ns) (Fig. 4A, Tables S3 and S5). In contrast, the FlAsH labeled sensors F1_smDA_ (1.15 ± 0.03 ns) and F1_dmDA_ (1.16 ± 0.02 ns) displayed a substantial decrease in the average CFP fluorescence lifetime compared to the wild-type sensor with 2.5 ± 0.02 ns (Fig. 4A, Table S3 and S5). At the same time, F1_smDA_ and F1_dmDA_ exhibited comparable E_FRET_ (∼63%), indicating that the mutations bring the donor and acceptor of the FRET pair into close spatial proximity which is drastically different from the F1_DA_ E_FRET_ of 19%. Next, we checked the effect of GephFL upon interaction with the mutant sensors. As expected, no average fluorescence lifetime change was observed in F1_smD0_ and F1_dmD0_ upon GephFL interaction (Table S5). However, to our surprise, an interaction of GephFL with F1_smDA_ and F1_dmDA_ resulted in a drastic increase in their average fluorescence lifetimes to 1.98 ± 0.03 ns and 2.05 ± 0.05 ns, respectively (Fig. 4A, Fig. S6), indicating a substantial increase in inter-fluorophore distance upon GephFL binding.

**Figure. 4.**
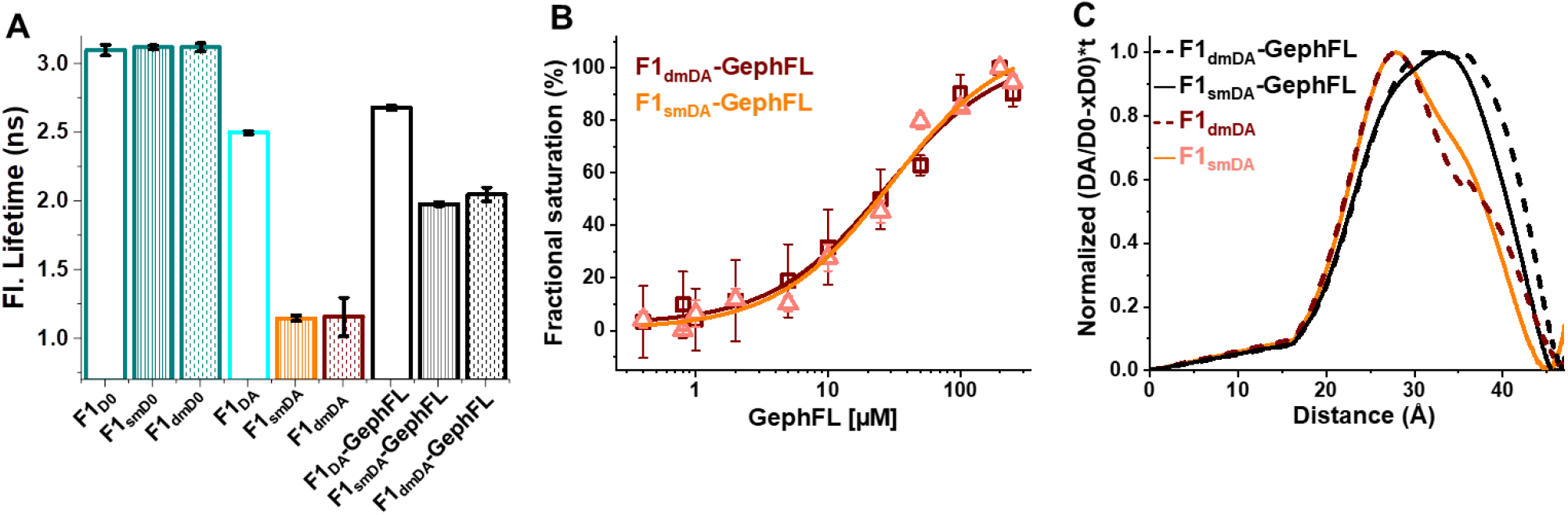
Mutant sensors construction and characterization. **(A)** Bar graph showing the species-weighted average fluorescence-lifetime of the CB wild-type, single mutant (sm) and double mutant (dm) FRET sensors prior to (F1_D0_, F1_smD0_ and F1_dmD0_), after FlAsH labeling (F1_DA_, F1_smDA_ and F1_dmDA_) and the FlAsH labeled sensors in the presence of a 100-fold molar excess of GephFL. **(B)** GephFL binding affinity plot of F1_smDA_ (orange) and F1_dmDA_ (wine). Binding affinities were determined by first converting ⟨τ⟩ into the fractional saturation using eq. 4. and the data were further fitted with eq. 5. The GephFL binding affinity constant (K_D_) for F1_smDA_ and F1_dmDA_ were measured as 29.3 ± 6.5 µM and 26.6 ± 5.7 µM, respectively. Data from three different batches of experiments are presented as mean values ± SD. Model-free description of the inter-fluorophore distance distribution underlying the time-resolved fluorescence intensities (eq. 12-13). Normalized distance distribution curves shown for F1_smDA_ and F1_dmDA_ in the absence and presence of GephFL. F1_smDA_ and F1_dmDA_ show a major peak at 28 Å and a shoulder at 36 Å, which is significantly different from F1_DA_, particularly for the major peak located in this case at ∼43 Å (Fig. S9D). Upon complexation with GephFL the major peak shifts to 36 Å with a weak shoulder at ∼28 Å.

We also investigated the GephFL affinity for the open state mutant sensors and hence separately titrated F1_smDA_ and F1_dmDA_ with increasing concentrations of GephFL. GephFL titration with F1_smDA_ and F1_dmDA_ led a gradual increase in their ⟨τ⟩ followed by saturation (Fig. S6 C-D). As assessed from the binding affinity values (Fig. 4B) of F1_smDA_ (K_D_ = 29.3 ± 6.5 µM) and F1_dmDA_ (K_D_ = 26.6 ± 5.7 µM), both exhibited a comparable and moderately strong binding affinity for GephFL. Interestingly, the GephFL binding affinity for the open state sensors was comparable to that of the wild-type sensor (F1_DA_). Our studies with the mutant sensors corroborate previous data (Soykan et al., 2014), which suggest that the disruption of the intramolecular interaction leads to a conformational switch within CB.

We also tried to analyze the fluorescent lifetimes with a Gaussian distance distribution model (eq. 11), however, the fast exponential decay of the fluorescent intensities in the beginning for both F1_smDA_ and F1_dmDA_ sensors made the fitting with the Gaussian distance distribution model challenging. Thus, we followed a model-free approach reported earlier (Peulen et al., 2017) to visualize the distance distribution underlying the time-resolved fluorescence intensities of F1_smDA_ and F1_dmDA_ (Fig. 4C, S7 and S8). For comparison, we analyzed the F1_DA_ and F1_DA_-GephFL time-resolved fluorescent intensities in the same way (Fig. S9). The distance distributions of F1_smDA_ and F1_dmDA_ revealed a main peak around 28 Å and a shoulder around 36 Å, indicating a disruption in the intra-molecular interaction between the SH3 and DH domains, which concomitantly became more flexible, in both the single and double mutant (Fig. 4C). Thus, the FlAsH present at the N-terminus and CFP at the C-terminus move closer to each other as reflected in the main peak at 28 Å (high FRET state) and a second peak around 36 Å is observed (low FRET state). Upon ligand interaction the F1_smDA_ sensor showed a shift of the main peak from 28 Å to 36 Å with a minor shoulder at 28 Å, further indicating that the distance between the donor and acceptor increases upon ligand interaction. A similar type of distance shift was also observed for F1_dmDA_ upon interaction with GephFL. In contrast, in case of F1_DA_ and the F1_DA_-GephFL complex (Fig. S9), the main peak is at ∼43 Å (low FRET) with a small shoulder at ∼25 Å (high FRET). Thus, the high FRET population is strongly increased in case of F1_smDA_ and F1_dmDA_ compared to F1_DA_ and the F1_DA_-GephFL complex, indicating that the opening of the structure due to mutations is different from the opening caused by ligand interaction of the wild type CBFRET sensor.

### Gephyrin binding elicits differential responses in a series of FRET sensors

To investigate the orientation of the SH3 domain with respect to the DH-PH tandem during activation, we performed interaction studies of GephFL with CB constructs displaying the tCM at three additional positions: After residue 28 and 73, i.e. at the start and end of the SH3 domain, assuming that these positions should be sensitive to SH3 domain motion during activation, and after residue 99, close to the DH domain, for understanding DH-PH domain movements. Initially, CFP ⟨τ⟩ measurements were carried out for the FlAsH-labeled sensors denoted as F28_DA_, F73_DA_ and F99_DA_. The highest ⟨τ⟩ decrease was observed for F73_DA_ (2.07 ± 0.01 ns) followed by F99_DA_ (2.28 ± 0.02 ns), whereas F28_DA_ (2.54 ± 0.01 ns) showed a similar value as F1_DA_ with 2.5 ± 0.01 ns (Fig. 5A and S10 A). Upon interaction with GephFL, F73_DA_ and F99_DA_ displayed a significant increase in ⟨τ⟩ with 2.4 ± 0.01 ns and 2.6 ± 0.01 ns (Fig. 5A and S10 B), respectively, whereas F28_DA_ (2.64 ± 0.01) displayed a comparable change in ⟨τ⟩ as observed for F1_DA_ (2.73 ± 0.02 ns) (Tables S3 and S6).

**Figure 5.**
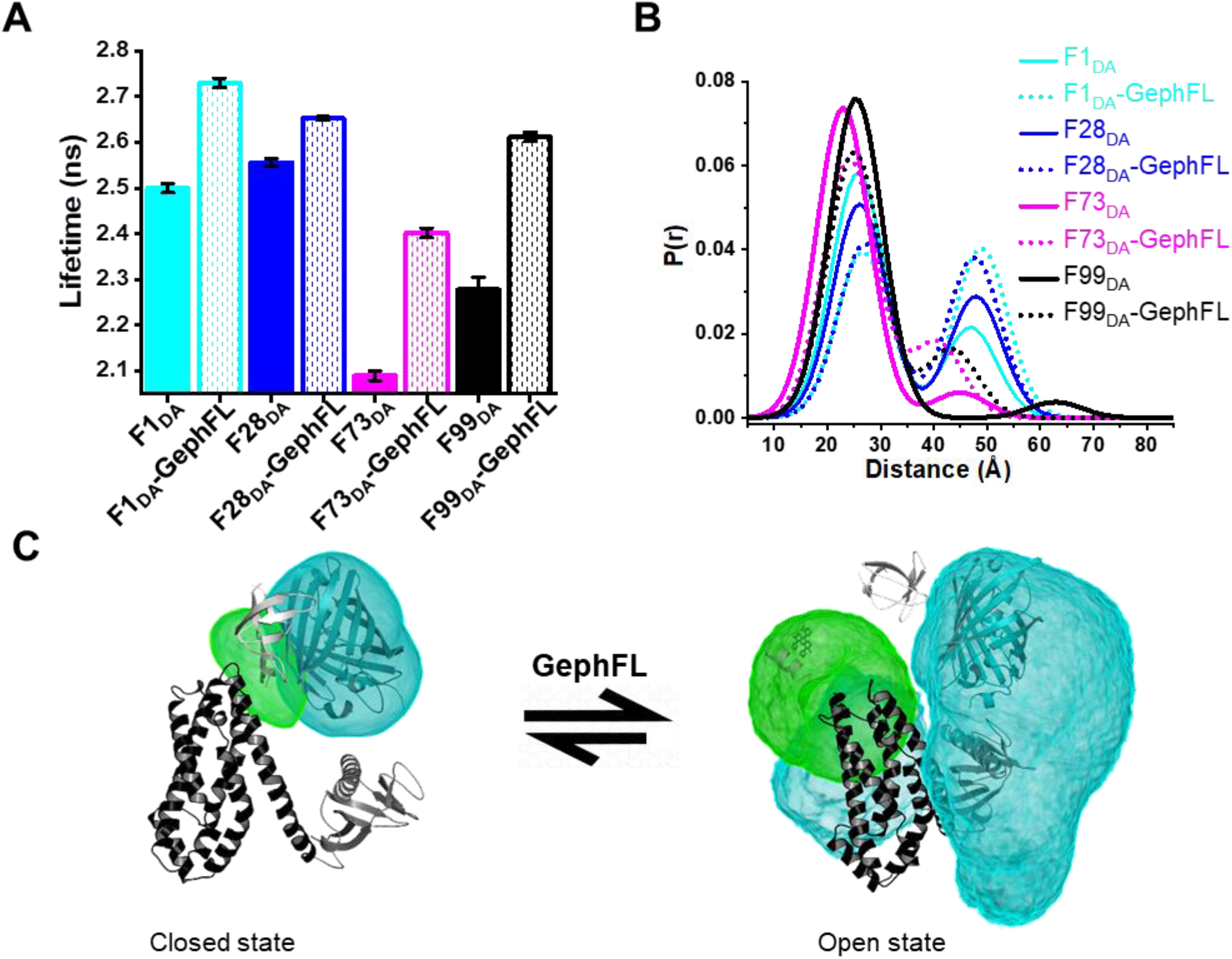
Characterization of additional CB FRET sensors and FRET-restraint based Markov-chain Monte Carlo (MCMC) sampling. **(A)** Average fluorescence lifetime observed of F1_DA_, F28_DA_, F73_DA_ and F99_DA_ in the absence (solid bars) and presence (dotted bars) of GephFL. **(B)** Distance distribution obtained from the two Gaussian distributed distances fit model of different FlAsH labeled sensors in the absence (solid lines) and presence (dotted lines) of GephFL. In case of F1_DA_ and F28_DA_ the change in the low FRET state upon interaction with GephFL is larger. **(C)** MCMC sampling of the fluorophore positions in CB based on the experimentally obtained FRET restraints (Table S8), connectivity restraints (Table S8) and the available structures for CB-SH3- (PDB ID 4mt7) and CB-SH3+ (PDB ID 4mt6). The cyan density reflects the spatially accessible volume of CFP, whereas the green color shows the spatially accessible volume of FlAsH incorporated at residue 99. The conformational change in CB from the free to the bound state allows both fluorophores to sample a much larger volume. In the bound state the excluded volume due to gephyrin binding should be considered.

Distance distribution studies for all sensors suggested that the absence or presence of GephFL does not alter the high-FRET and low-FRET states, indicating that all sensors including F1_DA_ displayed comparable inter-fluorophore distances (Fig. 5B, Table S6). The equilibrium between the high-FRET (x_1_) and low-FRET states (x_2_) upon GephFL interaction is comparable to F1_DA_ in case of F73_DA_ and F99_DA_ (Table S6). In contrast, F28_DA_ showed a comparatively smaller equilibrium shift following GephFL binding, as is evident from the fractions of the x_1_ and x_2_ species (Table S6). It must be noted that x_NoFRET_ is very low for F73_DA_ and F99_DA_ but increases significantly for all sensors upon interaction with GephFL (Table S6). In summary, the results clearly indicated that GephFL induces a movement of the SH3 and DH domain during the transition from the closed to the open conformation.

### Markov-chain Monte-Carlo sampling studies reveal CB conformations

To comprehend the overall conformational changes between the open and closed states of CB, especially with respect to SH3 domain, which can be described as “attached” and “detached” states, we performed Markov chain Monte-Carlo (MCMC) sampling studies based on the experimental distance distributions derived for our four FRET sensors (F1_DA_, F28_DA_, F73_DA_ and F99_DA_), their respective uncertainties (Table S7), and the molecular architecture of CB. As input for the MCMC sampling (Kalinin et al., 2012) the structures of CB without the SH3 domain (PDBID 4mt7) and with the SH3 domain in the closed state (PDBID 4mt6) were used and dissected into their domains connected by flexible hinges (see Methods, Table S8). For the sampling of the open, GephFL-bound state, the respective low FRET distances were used, while for the closed, GephFL-free state, the respective high FRET distances were used (Table S7). We evaluated the accessible volume of CFP, F99_DA_ (Fig. 5C and S12) and the SH3 domain in the open and closed state. As Fig. 5C shows, the opening and PH-domain rotation/tilt with respect to the DH-domain generates an “empty” space between those two domains and thus gives the C-terminally attached CFP and F99_DA_ access to a larger space, while in the closed state, both CFP and F99 are rather restricted and confined by the excluded volume of the DH-PH domains. In contrast, the experimental restraints for the closed state allow a relatively large accessible volume for the SH3 domain already in the closed state, on either side of the DH domain. The sampling of the open state for the SH3 domain resulted in a large, bilobed volume surrounding the DH-PH domains (Fig. S12 B), indicative of a less well resolved, freely diffusing SH3 domain. This large spatial offset of the SH3 domain with respect to DH-PH domain is in agreement with previous SAXS data (Soykan et al., 2014).

## Discussion

Despite the fundamental importance of CB as an adaptor protein ensuring the proper function of inhibitory GABAergic synapses, its interaction with the neuronal scaffolding protein gephyrin remained poorly understood. In the present study, we address this longstanding conundrum through the aid of custom designed CB FRET sensors. Here, we provide fluorescence lifetime-based FRET data, which, along with FRET-restrained modelling studies, elucidates the molecular mechanism of autoinhibition of CB and its activation by gephyrin. Previous studies hypothesized that activation of CB upon interacting with NL2_icd_ or Cdc42 leads to an open structure of CB, which allows CB to interact with phosphoinositides located in the postsynaptic membrane (Poulopoulos et al., 2009; Soykan et al., 2014). The two-state Gaussian distributed distance fit of the CB F1_DA_ sensor showed an increase in the population of the low FRET state upon interaction with NL2_icd_ indicating NL2_icd_-mediated CB opening. However, at comparable concentrations, NL2_icd_ displayed a smaller increase in average fluorescence lifetime as compared to GephFL, thus suggesting that GephFL not only interacts with but also efficiently activates CB. Compared to NL2_icd_ GephFL even possesses a higher capability for CB activation. Contrary to the previously hypothesized notion (Jedlicka et al., 2009; Soykan et al., 2014) our data suggest that initial CB relief by NL2 binding is not essential for gephyrin-CB complex formation. Through the aid of the F1_DA_ FRET sensor we could successfully quantify the previously unknown binding strength of the GephFL-CB complex, yielding a reasonably tight interaction (K_D_ = 20.2 ± 8.3 μM) between the two proteins (Fig. 2D). Furthermore, using a two-state Gaussian distributed distance fit model for F1_DA_, we demonstrate that CB can be described as a two-state system, encompassing a compact or high FRET state and a relaxed or low FRET state (Fig. S2B). Quantification of CB molecules in the high and low FRET state indicated that GephFL binding shifts the equilibrium from the closed state of CB (high-FRET F1_DA_) population towards the open (low-FRET F1_DA_) state (Fig. 2C). Additionally, the gradual increase of x_NoFRET_ upon addition of GephFL indicates that there might be another state beyond the measurable FRET distance of 49 Å for this FRET pair. Our FRET study therefore provides concrete evidence of GephFL-mediated CB opening, further strengthening the role of gephyrin as a CB activator. Identical affinities for GephLE (9.7 ± 2.7 µM) and GephE (9.4 ± 4.6 µM), along with their highly similar behavior in mediating the transition from closed to open states of CB (Fig. 3B-C), indicate that the E domain is responsible for CB activation. Additionally, a monomeric (dimerization-deficient) variant of the E domain (GephE_mm_) was also able to facilitate CB activation, which demonstrates that GephE dimerization is not a prerequisite for its interaction with CB. However, the low binding strength (69 ± 12 µM, Fig. 3B) of GephE_mm_ for F1_DA_ indicates that GephE dimerization is required to enhance its affinity for CB, potentially by stabilizing the E-domain.

The constitutively active mutant CB FRET F1_smDA_ and F1_dmDA_ sensors, somewhat counterintuitively, exhibited an increased average FRET efficiency compared to the wild-type F1_DA_ sensor. This might be because the disruption of the intramolecular SH3-DH/PH interactions rearranges the SH3 and PH domains and brings the FlAsH and CFP moieties into closer proximity. A comparable ⟨τ⟩ decrease (Table S5) in both mutant sensors along with their similar affinity for GephFL (Figure 4B), suggests that the single E262A variant is already capable of eliminating the intramolecular interactions between the SH3 domain and the DH-PH tandem and both mutant sensors potentially attain similar tertiary structures.

Structural insights into full-length CB is limited to a low resolution apo-CB-SH3^+^ crystal structure (Soykan et al., 2014) and hence information about the SH3 domain orientation in active CB state is limited. With our series of CB FRET sensors, we could visualize the accessible space of the SH3 domain in the active state of CB (Fig. 5C, Fig S12). In all sensors, GephFL addition led to an increase in the mean CFP fluorescence lifetime, thus suggesting an increase in the average inter-fluorophore distance in the respective sensors (Fig. 5A-B). Overall, the studies with the F28_DA_, F73_DA_ and F99_DA_ sensors provide strong evidence for a displacement of the SH3 domain following the interaction of CB with gephyrin. Furthermore, Markov-chain Monte-Carlo (MCMC) sampling (Greife et al., 2016; Kravets et al., 2016) based on the experimentally obtained inter-fluorophore distances of the four sensors and protein domain connectivity clearly identified distinct closed and open states of CB. In the closed conformation of F99_DA_ the two fluorophores were found to be in close spatial proximity. In contrast, GephFL addition led to a clearly distinguishable open state of CB, in which the probability densities of the two fluorophores were clearly separated (Fig. 5C, Fig. S12 A) with enhanced CFP density being present on the opposite side of the connecting helix between the DH and PH domains. This further suggests that GephFL-mediated CB opening causes a disruption of the intramolecular interaction between the SH3 domain and DH-PH tandem, thereby also rendering the PH domain flexible and generating space between the DH-PH domains, which can in turn be occupied by the fluorophores. Please note that gephyrin was not present in the MCMC modelling studies, however, it would significantly restrict the space accessible to the fluorophores. As is evident from the extended density of the SH3 domain in the closed state and the presence of both low and high FRET states in the gephyrin-free measurement, our lifetime-based FRET experiments and modelling studies suggest an equilibrium between an SH3-attached and SH3-detached state already in the inactive, unbound state (Fig. S12 B). This was not apparent in the available structure of the closed state where crystal packing forces presumably selected for a single closed state. However, only a flexibly attached SH3 domain, in equilibrium between an attached and detached state, would allow other proteins to bind in the region usually occupied by the SH3 domain.

Our *in vitro* and modelling data with CB FRET sensors led us to formulate a model (Fig. 6) summarizing the gephyrin-mediated CB activation. Taken together, our results reveal a clear interaction between full-length gephyrin (and its domain variants) and CB, an interaction that has been first identified more than 20 years ago (Kins et al., 2000), however, owing to technical limitations, could not be comprehensively characterized. Our data provide a framework to understand how CB acts as a dynamic molecular switch cycling between closed/inactive and open/active states in response to gephyrin binding.

**Figure 6.**
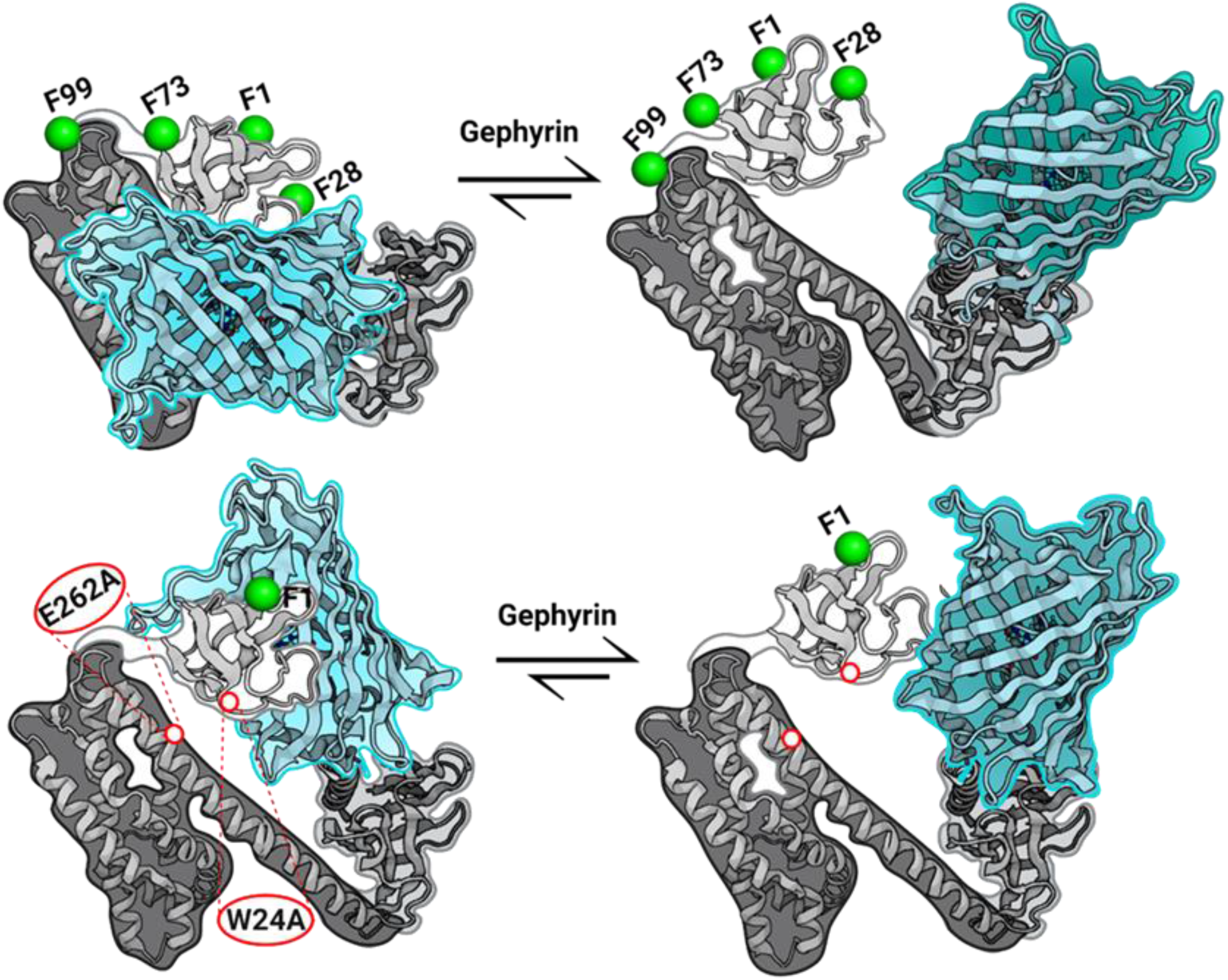
Schematic representation of gephyrin-mediated CB opening and complex assembly. **(A)** Cartoon and surface representation of full-length CB (PDBID 4mt6) in its auto-inhibited conformation depicting the individual domains (SH3; light grey with white surface, DH; black with dark grey surface and PH; grey with light grey surface). The figure illustrates the ensemble of sensors used in this study with the FlAsH attachment sites after amino acid residue 1 (F1_DA_), 28 (F28_DA_), 73 (F73_DA_) and 99 (F99_DA_) represented as green spheres, whereas CFP (cyan) was inserted after residue 456. Addition of gephyrin shifts the equilibrium towards an open state. In the closed, high FRET state, the C-terminally attached CFP exhibits significant quenching (cyan), whereas the low FRET, open state (CFP in teal) is characterized by reduced CFP quenching. **(B)** Cartoon representing the active state double mutant CB FRET sensor (F1_dmDA_, the mutated residues are indicated by white dots), which is already in an open state, and its conformational change upon interaction with gephyrin.

## Methods

### Cloning and Expression

The wild-type CB FRET sensor (F1_D0_) was constructed by inserting a tetra-cysteine motif (Adams et al., 2002) (tCM) after the first amino acid residue of the rat CB2-SH3^+^ splice variant, while CFP (Heim et al., 1994) was C-terminally attached (Fig. 1B, Fig. S1A) using restriction free (RF) cloning (Bond & Naus, 2012) in pETM14 (Table S2). Amino acid replacements for the open state mutant sensors (F1_smD0_ and F1_dmD0_) were generated by site directed mutagenesis. Additional sensors were created by inserting tCM at the specified position (Fig. 1B, Table S1). Full-length gephyrin (GephFL) and domain variants (GephG, GephE and GephLE) were previously described (Maric et al., 2014; Saiyed et al., 2007; Sander et al., 2013). For the GephE monomeric mutant (GephE_mm_) residues 318-750 were subcloned into the IMPACT system vector pTYB12 (New England Biolabs) and point mutations were introduced by site-directed mutagenesis. The SH3 domain of CB was generated by subcloning the cDNA coding for residues 10–79 (Soykan et al., 2014) of rat CB2-SH3^+^ into the pETM14 vector. The intracellular cytosolic domain of NL2 (NL2_icd_) encompassing residues 700-836 (Hoon et al., 2011) was subcloned into the pETM11 vector.

All CB FRET sensors were expressed in the *E. coli* strain BL21 (DE3). Cell lysates were subjected to affinity chromatography on Protino Ni-IDA Resin (Macherey Nagel) equilibrated in buffer A (50 mM Tris-HCl pH 8, 250 mM NaCl, 5% glycerol and 5 mM β-mercaptoethanol). Samples were eluted using buffer A containing 300 mM imidazole and were subsequently applied to a MonoQ 10/100GL column (GE Healthcare) and eluted using a linear NaCl gradient from 50 mM to 1 M NaCl. Finally, sensors were subjected to size exclusion chromatography on a Superdex 200 column (GE Healthcare) and concentrated by ultrafiltration. Full-length gephyrin and its domain variants (GephG and GephLE) were purified as described before (Sander et al., 2013) with small modifications. GephE and its dimer-deficient mutant (GephE_mm_) were initially subjected to affinity chromatography on chitin agarose beads (New England Biolabs), followed by ion exchange chromatography and subsequent size exclusion chromatography.

### Circular-dichroism spectroscopy

Circular-dichroism (CD) spectroscopy was performed with a Jasco J-810 spectropolarimeter. Prior to measurements, the buffer was exchanged to phosphate-buffered saline (PBS) pH 8.0 using ultrafiltration (Sartorius Vivaspin 500, Göttingen). Far-UV CD spectra from 190 to 260 nm were recorded at a scanning speed of 50 nm/min with a response time of 1 second and a bandwidth of 1 nm. CD spectra were recorded repeatedly (n=15) for each sample and averaged to optimize the signal to noise ratio. The buffer spectrum was also recorded and subtracted from the protein spectra. All measurements were conducted at room temperature.

### *In vitro* FLAsH Labeling

For FlAsH labeling (Adams et al., 2002; Griffin et al., 1998), purified sensors designated with the subscript D0 were first incubated at room temperature (RT) in labeling buffer (50 mM Tris-HCl pH 8, 250 mM NaCl, and 5 mM β-mercaptoethanol). Afterwards, FlAsH reagent (Cayman Chemicals) was added in a 10-fold molar excess to the sensors and the mixture was further incubated for 30 minutes at RT. Later, the mixture was dialyzed thoroughly against labeling buffer to remove unbound FlAsH, resulting in the corresponding DA fluorophore pair. Labeled sensors were flash frozen and stored at -80° C for later use.

### Time-resolved setup and data acquisition

Time resolved fluorescent measurements were conducted with a custom-built confocal microscope setup (IX 71, Olympus, Hamburg, Germany) equipped with a time-correlated single photon counting (TCSPC) system (Hydraharp 400, Picoquant, Berlin, Germany) with data acquisition by the fluorescence lifetime correlation software SymPhoTime 64 (PicoQuant, Berlin, Germany). The excitation laser (440 nm pulsed laser LDH-D-C-440, Picoquant) was fiber coupled (Laser Combining Unit with polarization maintaining single mode fibre, PicoQuant, Berlin, Germany) and expanded to a diameter of 7 mm by a telescope to fill the back aperture of the objective (60x water immersion, NA 1.2, Olympus, Hamburg, Germany), thus creating a diffraction-limited focal spot. Before entering the objective lens, the laser polarization was adjusted by an achromatic half-wave plate (AHWP05M-600, Thorlabs, Bergkirchen, Germany). A beam splitter (HC458 rpc phase r uf1, AHF) guided the laser through the objective, epi-illuminating the sample. In the detection path a 100 µm pinhole (PNH-100, Newport, Darmstadt, Germany) rejected out of focus light before being projected on photon counting detectors (2x PMA Hybrid-40, Picoquant, Berlin, Germany) by a telescope in a 4f configuration (focal length of lenses: 60 mm, G063126000, Qioptiq, Rhyl, UK). The beam was split via a polarizing beam splitter cube (10FC16PB.3, Newport, Darmstadt, Germany) in parallel (VV, detector 1) and perpendicular emission (VH, detector 2) after the first lens of the telescope. Emission filters (band pass filter Brightline HC 480/40 AHF, Tübingen, Germany) rejected unspecific light in each detection path. The laser was operated in pulsed mode at 20 MHz and a laser power at the sample of ∼11 µW, the temporal resolution was set to 4 ps. Measurements with the CB FRET sensors (1 µM concentration) were performed on standard glass coverslips (Menzel-Gläser, Braunschweig, Germany; 24 × 40 mm, 1.5).

CB FRET sensors (F1_D0_/F1_DA_) were titrated with varying concentrations of gephyrin (or other ligands) in binding buffer (50 mM Tris-HCl pH 8, 250 mM NaCl, 10 mM EDTA and 5 mM β-mercaptoethanol) in a final sample volume of 20 µL. Prior to data acquisition, samples were incubated overnight at 4 °Cunder dark light conditions. Samples were excited at 440 nm and the donor emission between 460 and 500 nm was recorded. Donor only (D0) and buffer solutions were measured as control samples and for background corrections. The data were acquired at room temperature for 5-10 min depending on photon counts. To determine the instrument response function (IRF), a KI-saturated solution of 3 µM fluorescein in double distilled water was measured for 10-15 min. To determine the relative detection efficiency in the parallel to the perpendicular channel, i.e. the g-factor, a 1 µM solution of Coumarin 343 was measured. Samples were measured in technical triplicates to calculate average and standard deviations for each condition.

### Time-resolved fluorescence decay analysis

To accurately determine the inter-fluorophore distance distribution from the fluorescence intensity decays the magic angle intensity decay was determined based on the obtained g-factor from the VV and VH signals. Data were exported from the ptu Symphotime format into text files using the Jordi-tool of the Seidel-Software package (www.mpc.hhu.de/software/3-software-package-for-mfd-fcs-and-mfis). Here, in the text file VV and VH data are stacked as a single column. All data were exported in 16 ps bins, i.e. 4096 channels for each detector. Thus, the single column text file contains 8192 channels in total with the first 4096 channels corresponding to the VV and the next 4096 channels to the VH decay. With a given g-factor, the analysis was done in the Chisurf software (Peulen et al., 2017) (https://github.com/Fluorescence-Tools/chisurf) as described elsewhere (Sanabria et al., 2020). We have calculated the g-factor to 0.9 from the tail fitting of the calibration dye coumarin 343. The decay curves were fitted with a multi-exponential model function using an iterative re-convolution approach (Sanabria et al., 2020; Tsytlonok et al., 2020) as follows

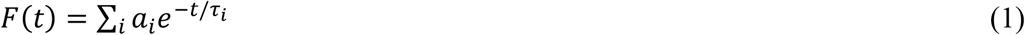

where *a*_*i*_ represents the amplitude and *τ*_*i*_ the lifetime of the corresponding component. Ideally, D0 should show a single component but, due to local quenching in donor-only samples, multi-exponential decays were expected. The quality of the curve fitting was evaluated by the reduced χ²-values and the weighted residuals. Time-resolved fluorescence intensities for FlAsH labeled (F1_DA_) and F1_DA_-ligand complexes (all gephyrin variants, NL2_icd_ and SH3) were also analyzed by eq. 1 to obtain the species-weighted average fluorescence lifetime.

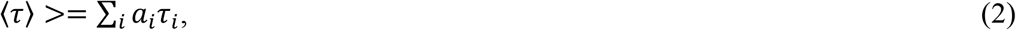

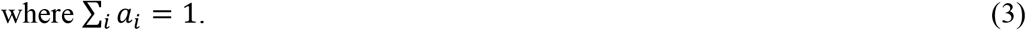

### K_D_ determination

We titrated the F1_DA_ sensor with different concentrations of full-length gephyrin (or other ligands) and measured the resulting time-resolved fluorescence intensities. The fractional saturation (in %) at concentration *i* was determined based on the average fluorescence lifetime:

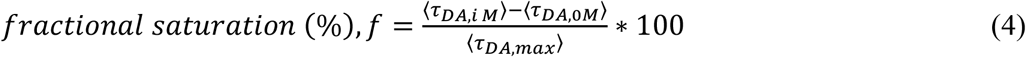

where ⟨*τ*_*DA,i M*_⟩ is the average fluorescence lifetime at concentration *i*, ⟨*τ*_*DA,0M*_⟩ is the mean fluorescence lifetime of the FlAsH labeled CB FRET sensor without addition of ligand and ⟨*τ*_*DA,max*_⟩ is the longest mean fluorescence lifetime of the titration, usually obtained at the highest ligand concentration. The resulting data points were plotted against the concentration of ligand and fitted as follows (Origin9, OriginLab):

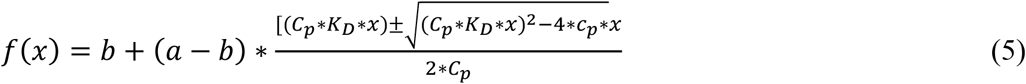

where *x* is the concentration, *b* the offset, *a*, the final intensity, *c*_*p*_ the protein concentration, and *K*_*D*_ the dissociation constant.

### Förster distance calculation

The Förster distance R0 [Å] was calculated from the overlap integral of the emission spectrum of the donor and absorption spectrum of the acceptor from

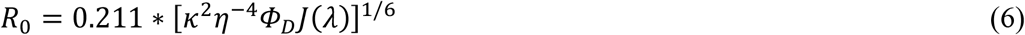

where κ^2^ is a factor describing the relative orientation in space of the transition dipoles of the donor and the acceptor. The magnitude of κ^2^ is assumed to be 0.66 for a random orientation of donor and acceptor. The refractive index (η) of the aqueous buffer is assumed to be 1.33. J(λ) is the overlap integral of emission of donor (CFP), and absorption (Fig. S1 C) of acceptor (FlAsH) and calculated by

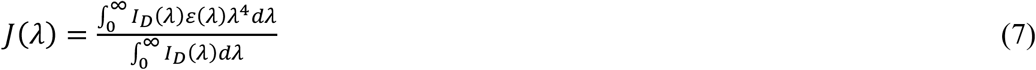

where I_D_(λ) is the fluorescence emission of the donor in the wavelength region λ and ε(λ) the extinction coefficient [M^−1^ cm^−1^] of the acceptor FlAsH (41000 M^-1^ cm^-1^ at 508 nm).

### Average FRET efficiency calculation

The fluorescence lifetime values obtained from the TCSPC decays were used to calculate an average FRET efficiency (E_FRET_) using the following equation:

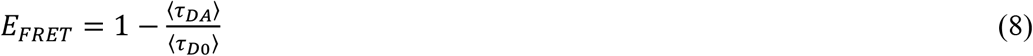

where ⟨*τ*_*D0*_⟩ and ⟨*τ*_*DA*_⟩ are the species-weighted average fluorescence lifetimes in the absence (D0) and presence (DA) of FlAsH as calculated based on eq. 2.

### FRET distance distribution analysis

For distance distribution analysis for the FlAsH labeled (F1_DA1_) and F1_DA1_-ligand complexes we followed a method described earlier (Sanabria et al., 2020; Tsytlonok et al., 2020). The time-resolved fluorescence intensities of the FRET-sample and the donor-only reference sample are presented as:

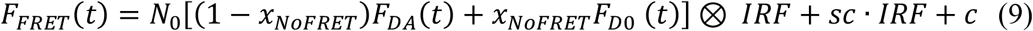

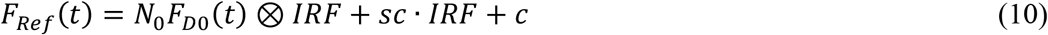

Here, *sc* is the scattered light from the sample, *c* is the constant offset of the fluorescence intensity and *N*_*0*_ is the total photon number. *x*_*NoFRET*_ is the no-FRET contribution from the unquenched donor. As stated earlier, we obtained multi-exponential fitting for the donor-only sample due to local quenching, however, the local quenching of the donor is not affected by FRET (Lehmann et al., 2020). Hence, the FRET-rate (*K*_*FRET*_) depends on the relative orientation and donor-acceptor-distance and the FRET samples can be fitted globally with the donor-only reference sample. In the presence of FRET, the donor fluorescence decay can be expressed with a Gaussian distance distribution (*ρ*) of donor-acceptor as

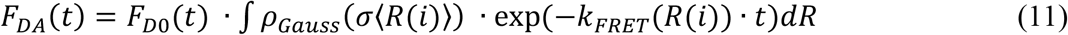

where ⟨*R*(*i*)⟩ is the mean distance between donor and acceptor and σ the width of the inter-fluorophore distance distribution *R*(*i*). In the analysis, σ was fixed to a physically meaningful value of 5 Å (Peulen et al., 2017). The Förster radius for CFP and FlAsH was 39 Å, as calculated following the method described in supplementary materials.

### Uncertainty estimation of distance distribution

The experimental uncertainty in the TCSPC-based inter-fluorophore distance analysis mainly stems from three sources: (i) The uncertainty of the orientation factor ⟨κ^2^⟩, δR_DA⟨κ2⟩_, (ii) the uncertainty in the D_only_ reference δR_DA_,reference (based on sample preparation etc.) and (iii) the statistical uncertainty based on the fitting, δR_DA_,fit (Peulen et al., 2017). Here, we estimated the uncertainty δR_DA_,fit in the obtained distances (Gaussian distance distribution, eq. 11) by sampling the *χr*^2^-surface in 50 steps in the range from -20% to + 20% of the respective distance using the “Parameter Scan” option in ChiSurf (Peulen et al., 2017). The value of the scanned distance, R_1_ or R_2_, respectively, is fixed, all other parameters are fitted and the resulting *χr*^2^ is reported. The resulting *χr*^2^-surface (Lakowicz, 2013) was plotted against the scanned distance (Fig. S11B) and the limits were determined using a 3σ-criterion based on an F-test (1700 TCSPC channels, 9 parameters) to a relative *χr*, ^2^ = *χr*,_i_ ^2^/*χr*,_min_ ^2^ of 1.012. To incorporate δR_DA,reference_ we extended the limits for R_min_ and R_max_ in such a way that the overall R_min_ and R_max_ for the experimental triplicates were used (Fig. S11B). The uncertainty of the orientation factor ⟨κ^2^⟩, δR_DA⟨κ2⟩_, which is usually the largest source of uncertainty, was not considered.

### Model-free distance distribution analysis

For a model-free description, we calculated the FRET-induced donor decay as described (Peulen et al., 2017). Briefly, in a first step, the fluorescence decay of the FRET sample *I*_*DA*_(*t*) is divided by the (modeled) decay of the single-labeled sample *I*_*D0*_(*t*). Next, the DOnly fraction, *x*_*NoFRET*_, i.e. the offset, is subtracted, and finally, this ratio is multiplied with the time axis *t* to yield the FRET-induced donor decay *ε*(*t*):

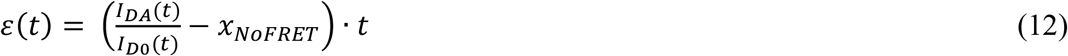

For an intuitive display, we converted the x-axis from time *t* to critical distance *R*_*DA,c*_ by the following relation:

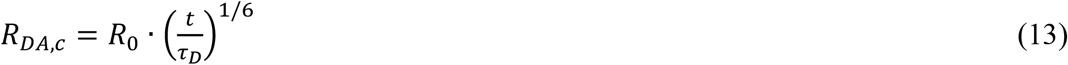

where *R*_*0*_ is the Förster radius of the respective FRET dye pair (here 39 Å) and *τ*_*D*_ the reference fluorescence lifetime of the donor fluorophore (here, 3 ns). Plotting *ε*(*t*) against *R*_*DA,c*_ results in a peaking distribution, which reflects the probability density function of the underlying distance distribution of the original decay *I*_*DA*_(*t*).

### FRET-restrained Markov-chain Monte-Carlo sampling

FRET-restrained Markov-chain Monte-Carlo (MCMC) sampling was performed using the FRET Positioning Software (FPS) (Kalinin et al., 2012) based on the X-ray structures of the CB-SH3^-^ (PDBID 4mt7) and CB-SH3^+^ (PDBID 4mt6) structures (Soykan et al., 2014). Additionally, the NMR structure of the ReAsH-tag (BioMagResBank ID code 16041) (Madani et al., 2009) was used as a model for the FlAsH-tag and the X-ray crystal structure of eGFP served as a template for CFP (PDBID 4eul). Two types of restraints were defined: (a) Connectivity restraints and (b) FRET-based restraints (Fig. S11A).

Connectivity restraints are based on the linear connectivity between the protein domains and labels, i.e. the SH3-domain is separated from the DH domain by 35 amino acids based on the structural models and 28 residues from the FlAsH-label incorporated at position 99 (Table S8). These flexible connections are modelled as worm-like chain polymers, where the uncertainty (or width of distribution) for FPS was determined as the 1σ-region. One exception is the connection between the PH domain and the CFP. Here, we assumed that residues 439-456 stay in their α-helical conformation as found in PDB entry 2dfk (Xiang et al., 2006), while residue 439 serves as a flexible hinge around which the helix and the ensuing CFP move about. Here, the uncertainty was set to the length of one amino acid residue (3.6 Å).

FRET-based restraints were based on the experimental distances obtained by the Gaussian distance fitting. The mean distance was set as the average from the experimental triplicates and the uncertainties were determined as described above (see Table S7).

Next, the five entities (FlAsH-1, SH3-domain (from 4mt6), FlAsH-99, CB (4mt6 or 4mt7) and CFP) were loaded into FPS and the respective CB structure (4mt6 or 4mt7) was fixed in place. The structural models were docked for 100 times, followed by 100x sampling using MCMC of each generated structure. In this step, the reciprocal kT was lowered to a value of 2. The resulting 10,000 models were exported as pml files, translated into PDB format and the docking and sampling results were verified by comparing the obtained restrained mean value with our input values. Next, the trajectories were generated using mdtraj (Robert et al., 2015) and the density based on the occupancy of the FlAsH-1, FlAsH-99, CFP and SH3-domain was exported from VMD (Humphrey et al., 1996). The data were visualized using PyMol (The PyMOL Molecular Graphics System, Schrödinger, LLC).

### Statistical analysis

All quantitative data are expressed as mean values ± standard deviation (SD) unless stated otherwise. Origin 9 (OriginLab) was used for statistical analysis. One-way ANOVA followed by Tukey’s post hoc multiple comparison test was performed for comparison between multiple pairs.

## Supporting information

This article contains supporting information.

## Data availability

All data needed to evaluate the conclusions in the paper are present in the paper or the supplementary materials. Raw data are available from the corresponding authors upon request.

## Author contributions

H.S. and K.G.H. Conceptualization; N.I., S.C. and K.H. data curation; N.I., S.C. and K.H. formal analysis; N.I., S.C. and K.H. Methodology; N.I., S.C., K.H., K.G.H. and H.S. project administration; H.S. and K.G.H. Supervision; NI and HS Writing-original draft; HS, NI, SC, KH and KGH Writing-review and editing; HS and KGH funding acquisition.

## Acknowledgements

We would like to thank the Seidel Lab (Molecular Physical Chemistry, Heinrich-Heine-Universität, Düsseldorf, Germany) for providing the MFD software package and Nicole Bader for excellent technical assistance.

## Abbreviations

The abbreviations used are

GABA,: aminobutyric acid
FRET, Fö: rster resonance energy transfer
NL2: Neuroligin-2
Geph: Gephyrin
CB: Collybistin
PH: pleckstrin homology
GEF: guanine nucleotide exchange factor.

## Results

