## Supporting Information for "Deciphering the conformational dynamics of gephyrin-mediated collybistin activation"

or

Katrin G. Heinze

† These authors contributed equally to this work

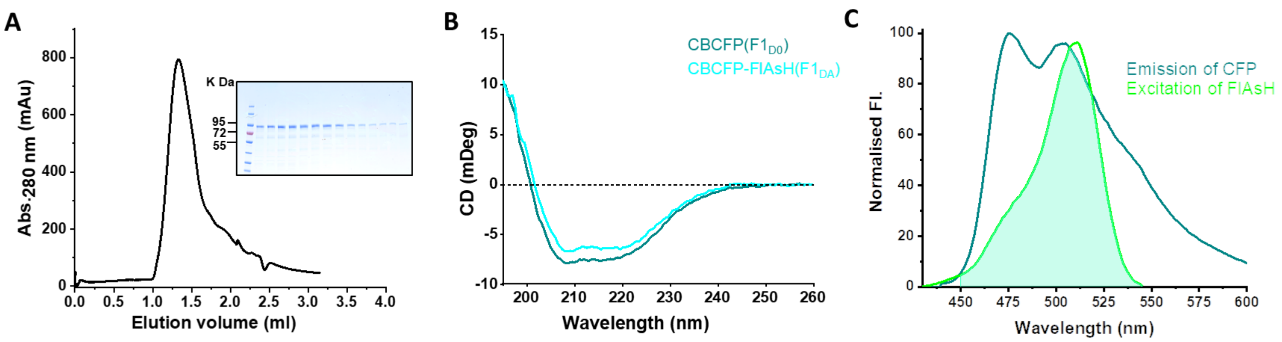


**Figure S1. CB FRET sensor purification and characterization.** (A) Elution profile of the CB $F1_{D0}$ FRET sensor (black) and corresponding SDS-PAGE gel showing the eluted protein from the size exclusion chromatography. (B) CD spectra of $F1_{D0}$ (teal) and $F1_{DA}$ (cyan). (C) Emission and excitation spectra of $F1_{D0}$ (teal) and $F1_{DA}$ (green), respectively. The filled area marks the overlap integral used for determining the Förster radius $R_0$.

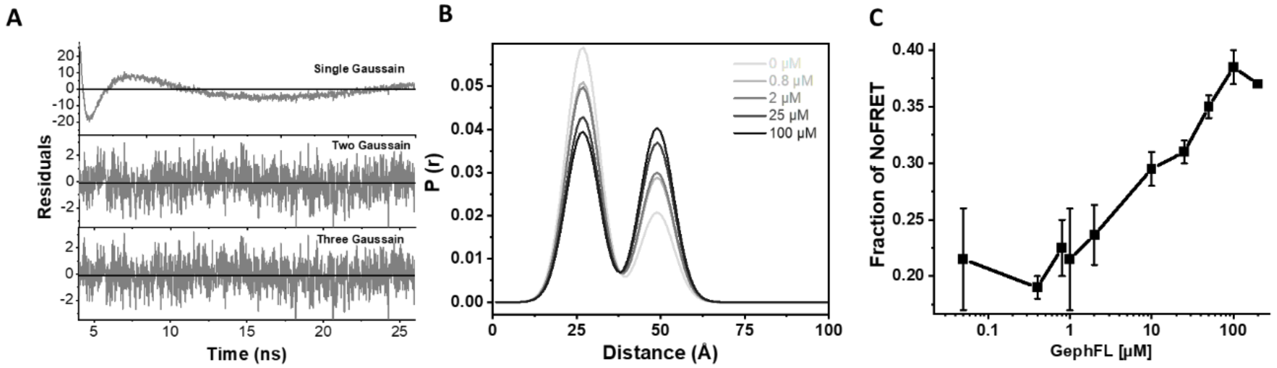


**Figure S2. Gaussian distributed distance fitting.** (A) Weighted residuals for fitting of the FlAsH-labeled CB FRET sensor ($F1_{DA}$) with one ($\chi_r^2$ = 47.14), two ($\chi_r^2$ = 1.14) or three ($\chi_r^2$ = 1.12) Gaussian distributed distances with a width of 5 Å. (B) The distance distribution based on the two state Gaussian distribution model shows that the fraction of the high FRET state decreases, and the fraction of the low FRET state increases upon increasing concentration of GephFL. (C) Increase of the No-FRET fraction, $x_{noFRET}$, of $F1_{DA}$-GephFL complex with increasing concentrations of GephFL, which may indicate the existence of an additional state exhibiting an inter-fluorophore distance >49Å where the FlAsH-CFP FRET pair is blind.

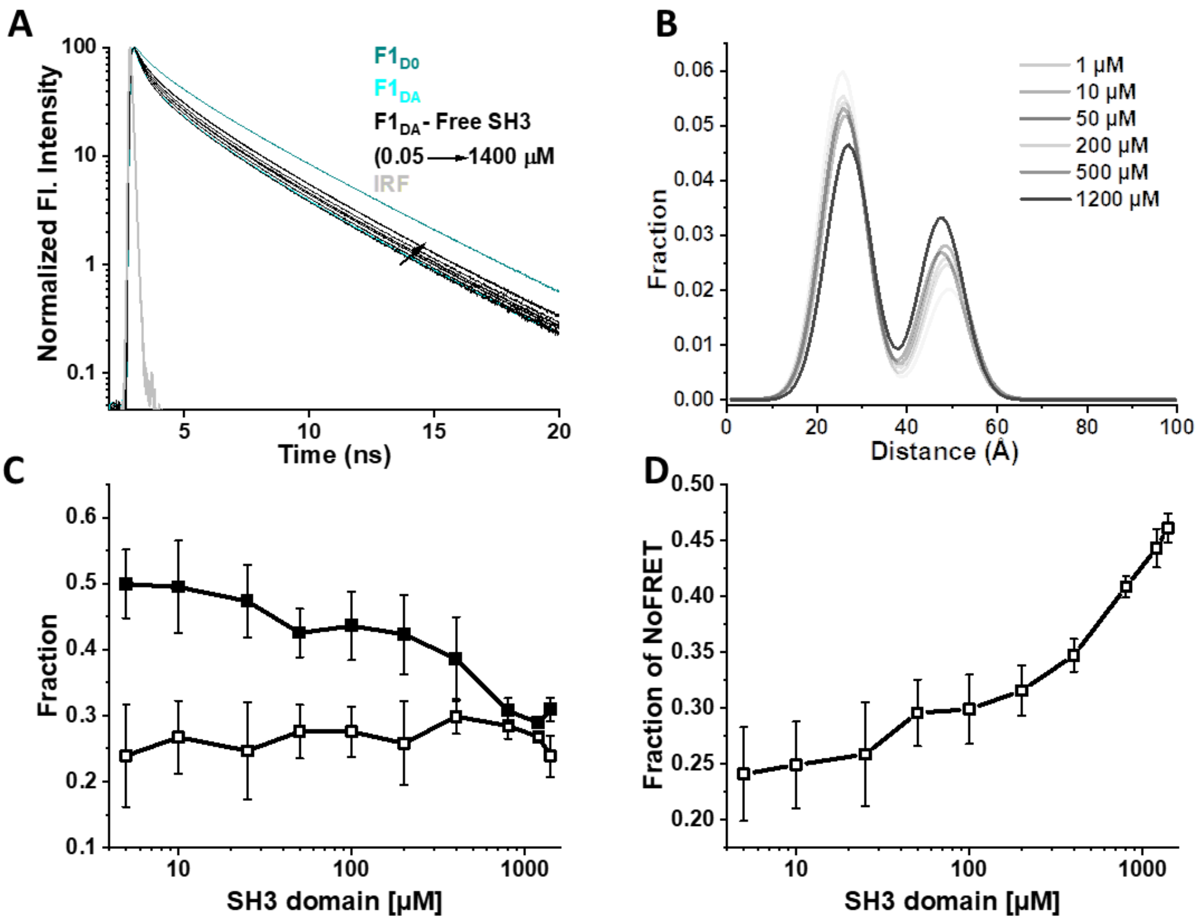


**Figure S3. Free-SH3 domain mediates CB opening.** (A) Time-resolved fluorescence intensities of the CB FRET sensor $F1_{D0}$ (teal), $F1_{DA}$ in the absence (cyan) and presence (black) of increasing concentrations of free-SH3 domain. Data were scaled to a maximum of 100 for easier comparison. (B) Distance distribution for the FlAsH labeled CB FRET sensor $F1_{DA}$ (cyan) with increasing concentrations of free SH3 domain. (C) Fraction of $F1_{DA}$ molecules in the closed/high FRET state (solid black squares) and their gradual transition into the open/low FRET state (unfilled black squares) upon addition of free SH3 domain. (D) $x_{noFRET}$ of the $F1_{DA}$-SH3 complex increases with increasing concentrations of GephFL. Again, this indicates another state with distances >49 Å.

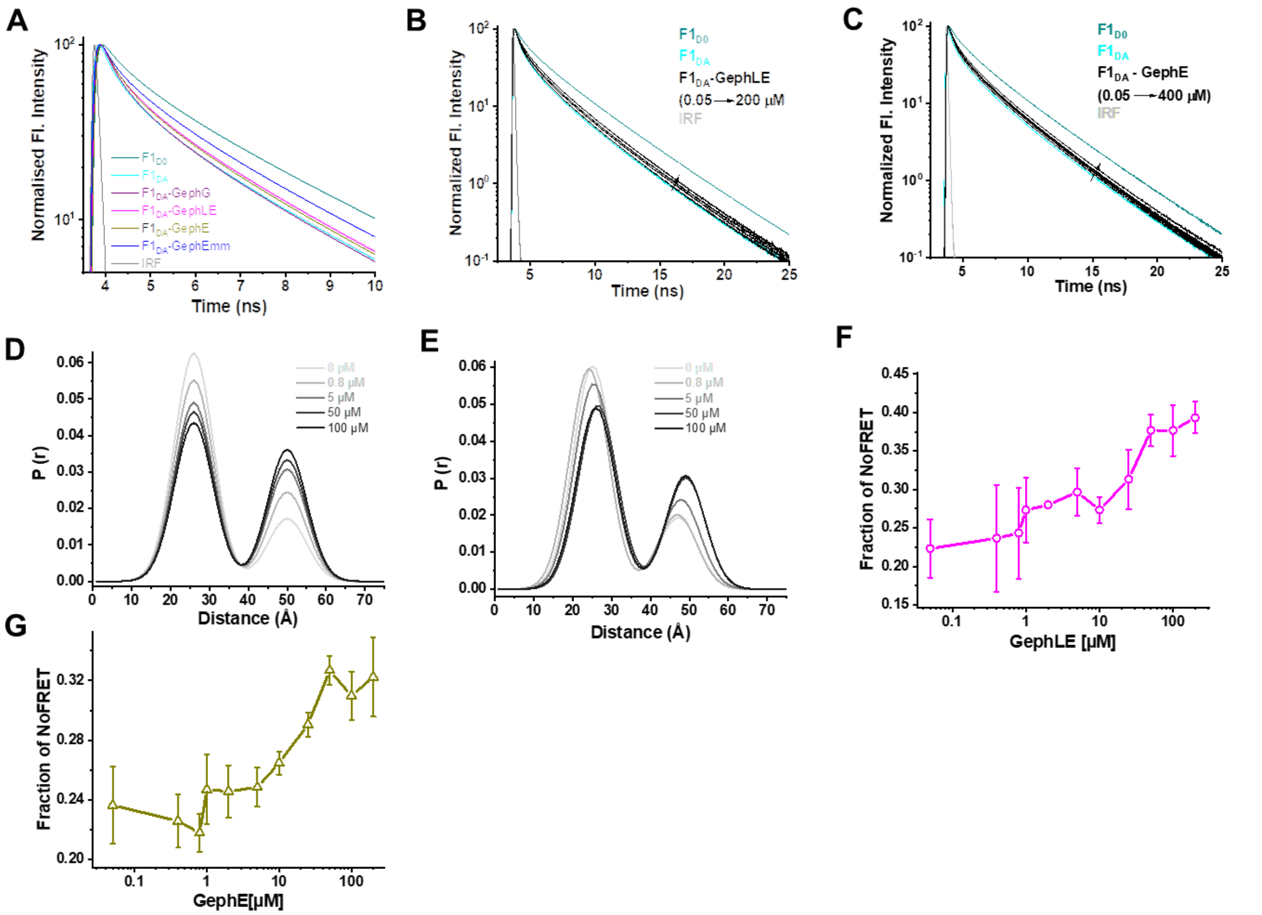


**Figure S4. Full-length gephyrin and variants containing the E domain facilitate CB opening.** (A) Time-resolved fluorescence intensities of CFP in $F1_{D0}$ (teal), $F1_{DA}$ (cyan) and in the presence of 100 µM GephG (purple), GephE (dark yellow) and GephLE (magenta) with IRF in light grey. Data were scaled to a maximum of 100 for easier comparison. (B-C) Fluorescence lifetime of the CB FRET sensor $F1_{D0}$ (teal), $F1_{DA}$ in the absence (cyan) and presence (black) of increasing molar concentrations of GephLE (B) and GephE domain (C), respectively. Data were scaled to a maximum of 100 for easier comparison. (D-E) Distance distribution of FlAsH labeled CB FRET ($F1_{DA}$) sensor upon interaction with various concentrations of GephLE/GephE. With increasing concentrations of GephLE/GephE the fraction of the high FRET state gradually decreases while the fraction of the low FRET state gradually increases. (F-G) Again, $x_{noFRET}$ of the CB FRET sensor increases upon interaction with GephLE/GephE.

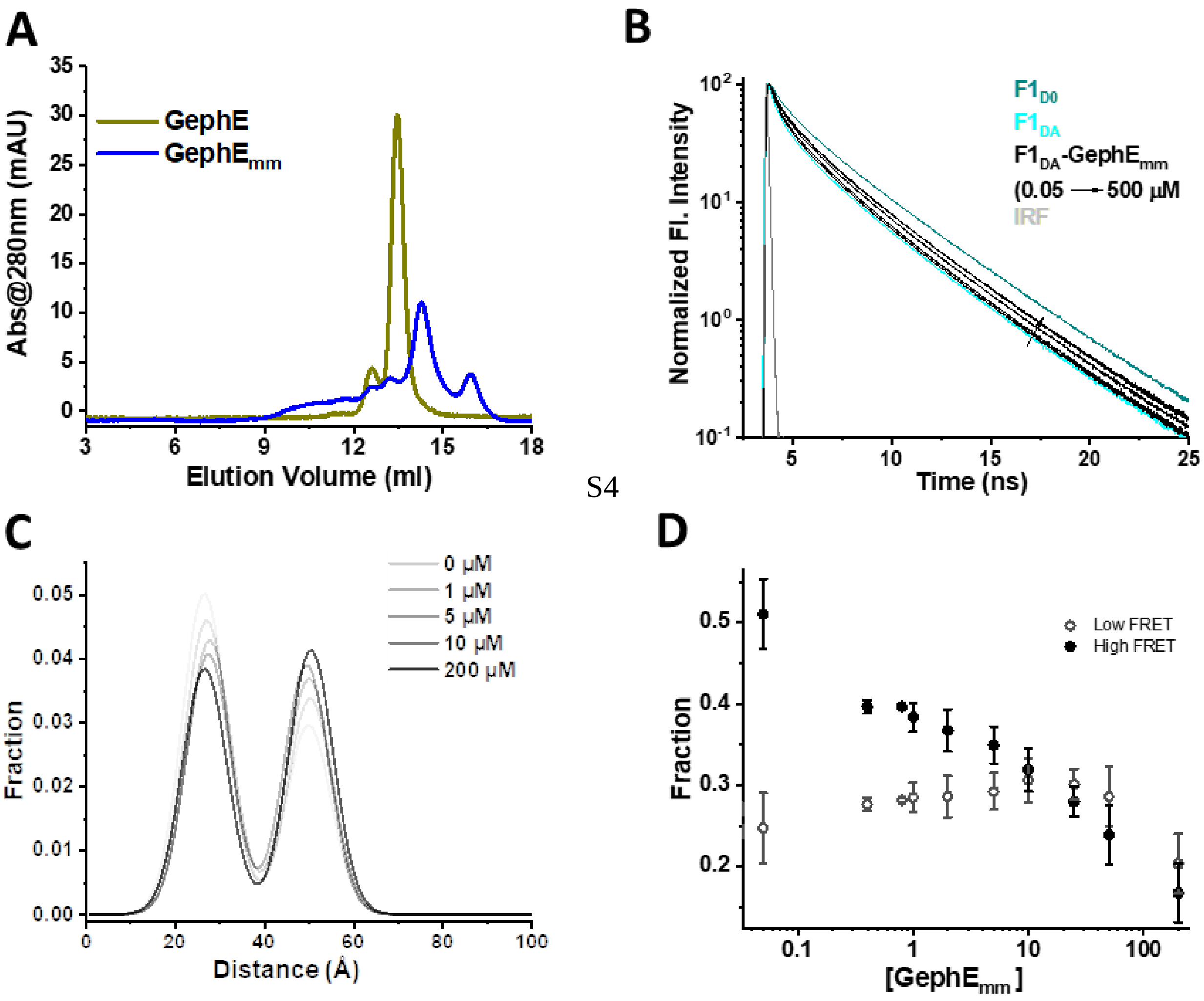


**Figure S5. GephE$_{mm}$ mediates CB opening.** (A) Size exclusion chromatography profile of GephE (yellow) and GephE$_{mm}$ (blue). (B) Time-resolved fluorescence intensities of CFP in F1$_{D0}$ (teal), F1$_{DA}$ alone (cyan) and in the presence of increasing concentrations of GephE$_{mm}$ (black). Data were scaled to a maximum of 100 for easier comparison. (C) Gaussian distance distribution analysis shows the decrease of the fraction of high FRET and concomitant increase in the fraction of the low FRET state with increasing concentrations of GephE$_{mm}$. (D) Fraction of F1$_{DA}$ molecules in the closed/high FRET state (filled circles, black) and their transition into the low FRET F1$_{DA}$ state (empty circle) upon addition of GephE$_{mm}$.

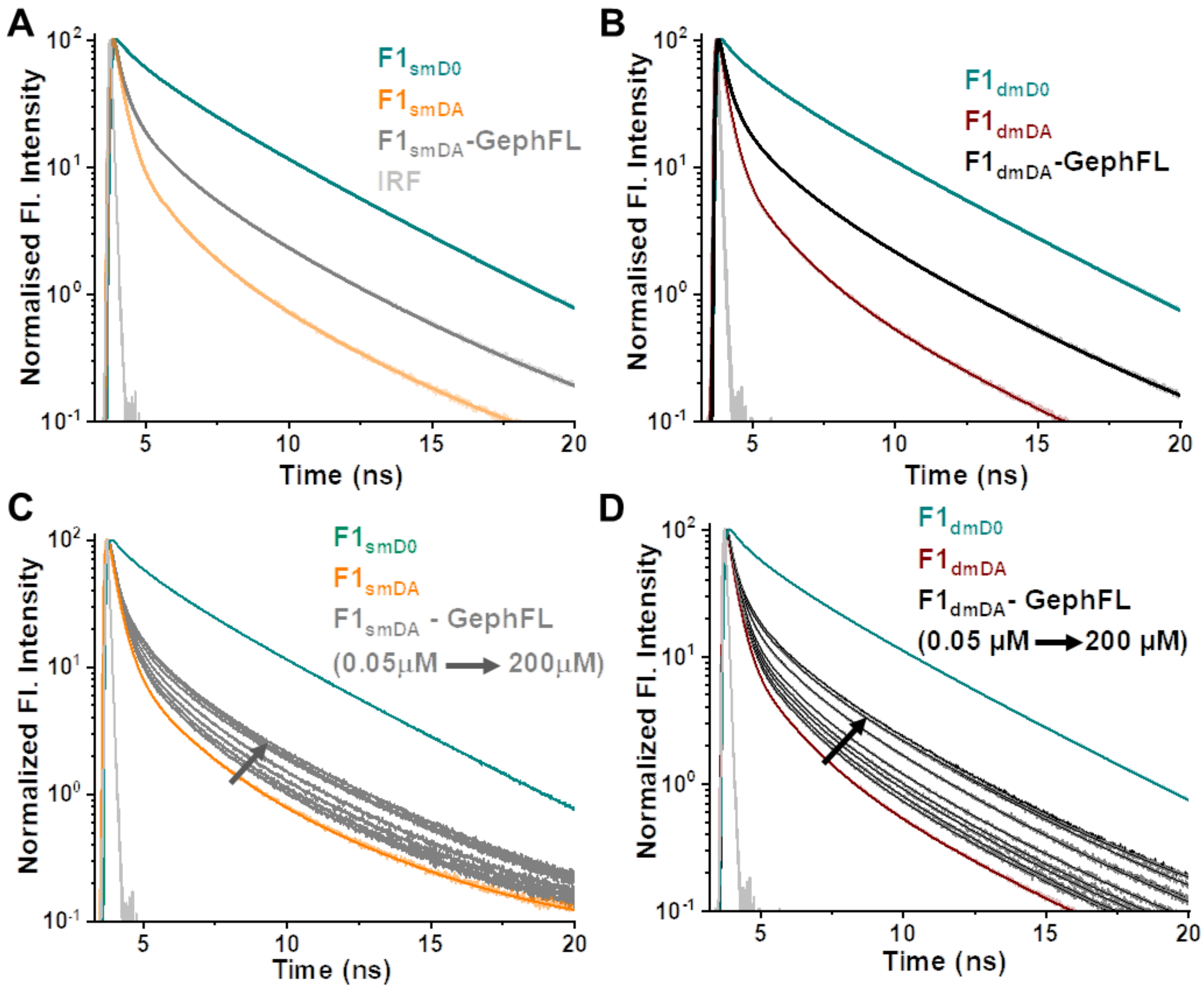


**Figure S6. Open state mutant sensor characterization.** (A) CFP fluorescence intensity decay of the single mutant FRET sensor $F1_{smD0}$ (teal) and the FlAsH labeled single mutant $F1_{smDA}$ sensor in the absence (light red) and presence (grey) of GephFL. (B) Fluorescence intensity decay of the double mutant FRET sensor $F1_{dmD0}$ (teal) and the FlAsH labeled double mutant FRET sensor $F1_{dmDA}$ in the absence and presence of GephFL in dark red and black, respectively. (C-D) Fluorescence intensity decays of $F1_{smDA}$ (C) and $F1_{dmDA}$ (D) in the presence of varying concentrations of GephFL. Data were scaled to a maximum of 100 for easier comparison. IRF is shown in light grey.

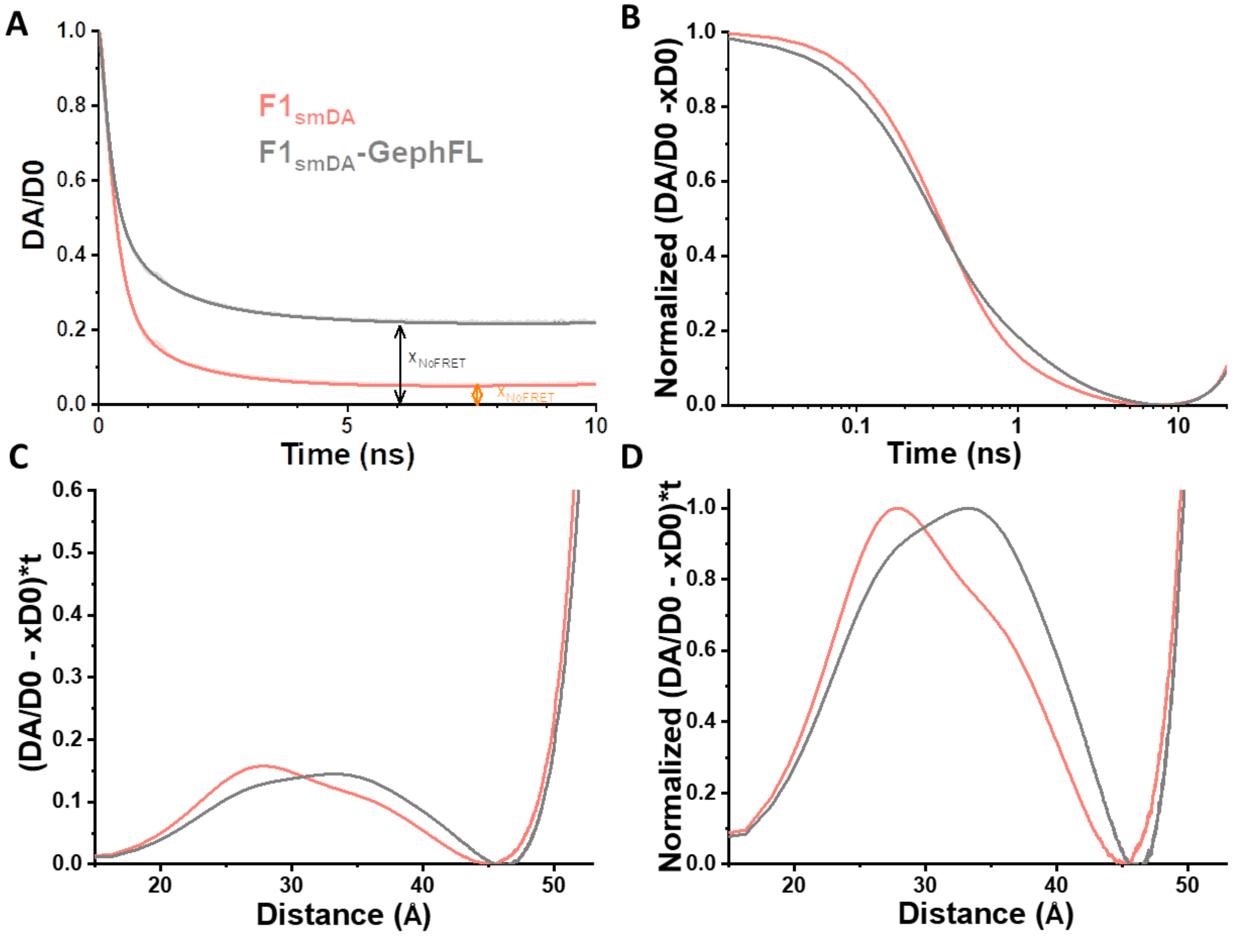


**Figure S7. Model-free visualization of the distance distribution underlying the time-resolved fluorescence intensities of $F1_{smDA}$ in the absence and presence of GephFL.** (A) The time-resolved fluorescence intensity of the double-labeled sample $I_{F1smDA}(t)$ is divided by the single-labeled sample $I_{F1smD0}(t)$. The offset values are the corresponding $x_{NoFRET}$ values (~0.05 for F1smDA and ~0.25 for F1smDA-GephFL). (B) The fraction of molecules not showing FRET ($x_{noFRET}$ or $x_{D0}$) – the constant offset in (A) – is subtracted and the time scale is logarithmic. (C) The time-axis is converted to the distance axis (eq. 11 main text). (D) The probability density distribution of the underlying distance distribution is normalized to 1 for easier comparison.

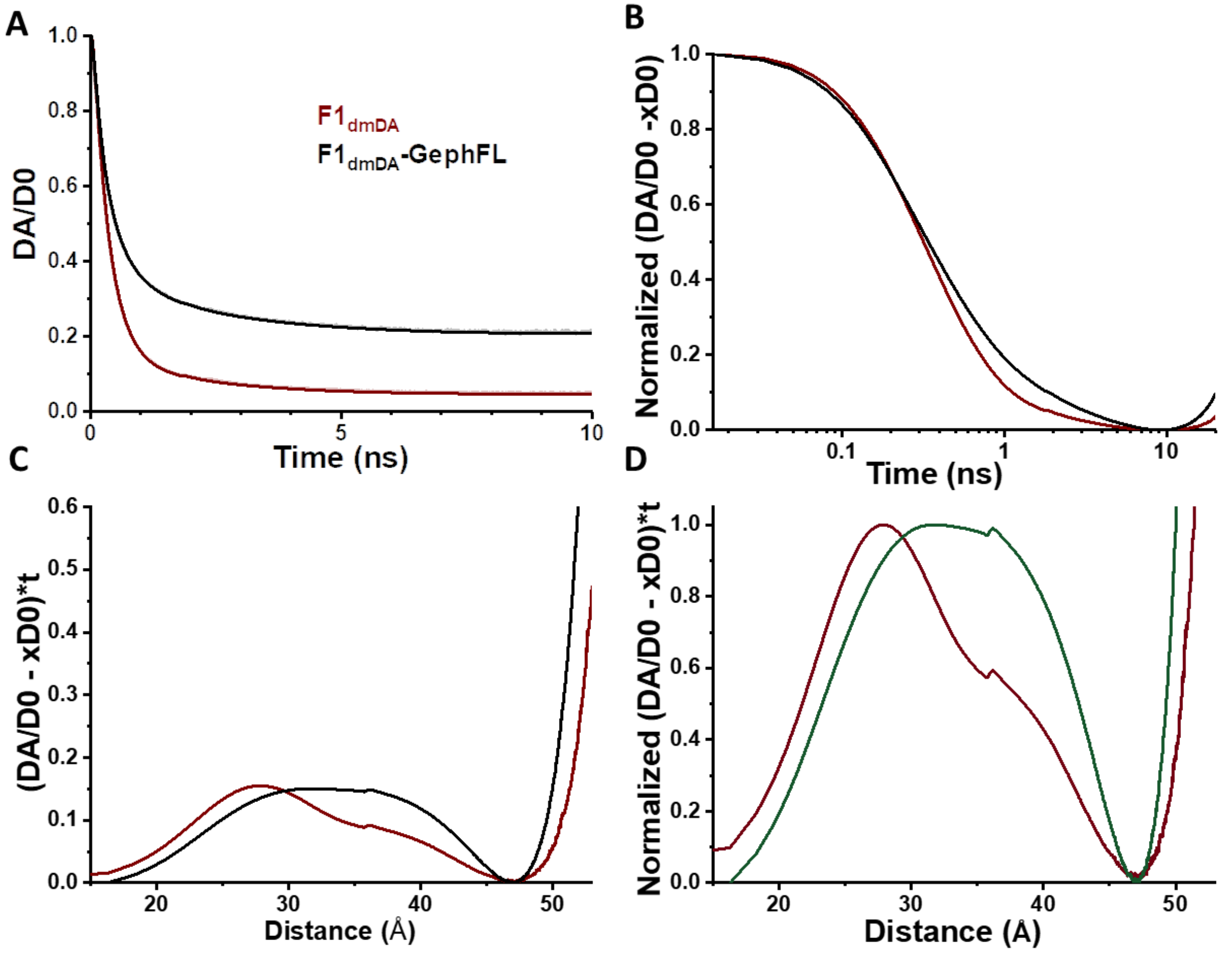


**Figure S8. Model-free visualization of the distance distribution underlying the time-resolved fluorescence intensities of $F1_{dmDA}$ in the absence and presence of GephFL.** (A) The time-resolved fluorescence intensity of the double-labeled sample $I_{F1smDA}(t)$ is divided by the single-labeled sample $I_{F1smD0}(t)$. (B) The fraction of molecules not showing FRET ($x_{noFRET}$ or $x_{D0}$) – the constant offset in (A) – is subtracted and the time scale is logarithmic. (C) The time-axis is converted to the distance axis (eq. 11 main text). (D) The probability density distribution of the underlying distance distribution is normalized to 1 for easier comparison.

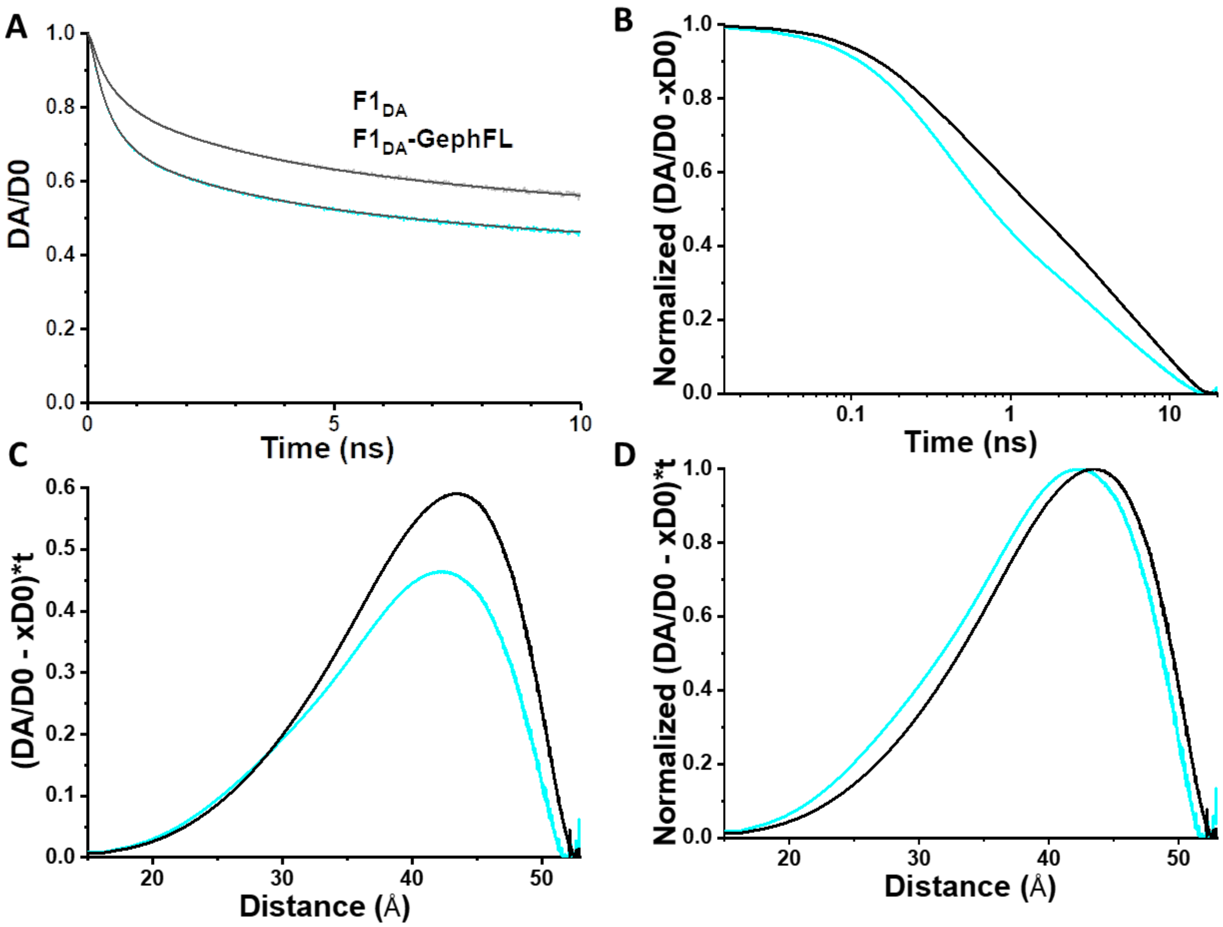


**Figure S9. Model-free visualization of the distance distribution underlying the time-resolved fluorescence intensities of F1$_{DA}$ in the absence and presence of GephFL.** (A) The time-resolved fluorescence intensity of the double-labeled sample $I_{F1smDA}(t)$ is divided by the single-labeled sample $I_{F1smD0}(t)$. (B) The fraction of molecules not showing FRET ($x_{noFRET}$ or $x_{D0}$) – the constant offset in (A) – is subtracted and the time scale is logarithmic. (C) The time-axis is converted to the distance axis (eq. 11 main text). (D) The probability density distribution of the underlying distance distribution is normalized to 1 for easier comparison.

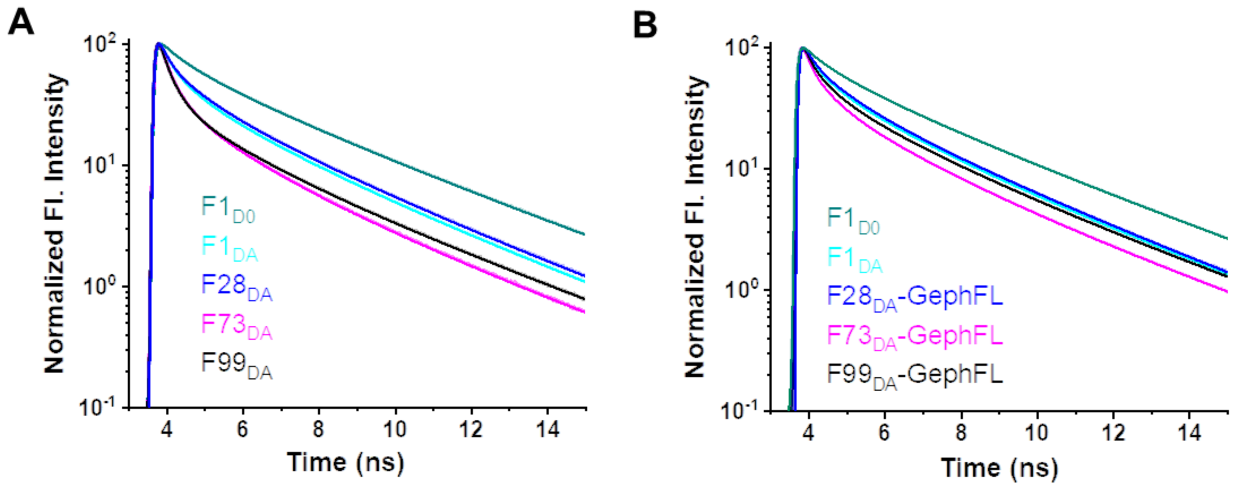


**Figure S10. Characterization of the additional CB FRET sensors.** (A) Comparative fluorescence intensity decay for the series of FlAsH labeled CB FRET sensors. Data were scaled to a maximum of 100 for easier comparison. Maximum quenching of the fluorescence lifetime was observed with $F73_{DA}$, whereas $F1_{DA}$ and $F28_{DA}$ show similarly low quenching. (B) Fluorescence intensity decays for the FlAsH labeled CB FRET sensors in the presence of full-length gephyrin. Data were scaled to a maximum of 100 in both figures for better comparison.

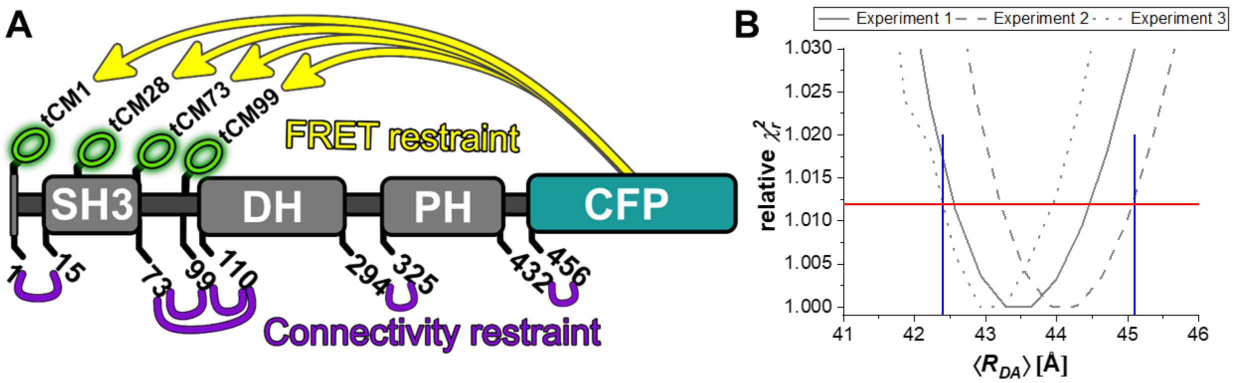


**Figure S11. FPS-based collybistin modelling.** (A) Connectivity and FRET-based restraints for collybistin modelling. Small numbers denote residue numbers based on the full-length collybistin sequence to which CFP is directly attached. (B) Exemplary $\chi^2_r$-surface for the $F1_{DA}$ – CFP low FRET distance of the open, GephFL-bound state. The red horizontal line indicates the 3σ-criterion of the $\chi_{r,rel}^2$ = 1.012, while blue vertical lines indicate the limits used for the FPS-based modelling.

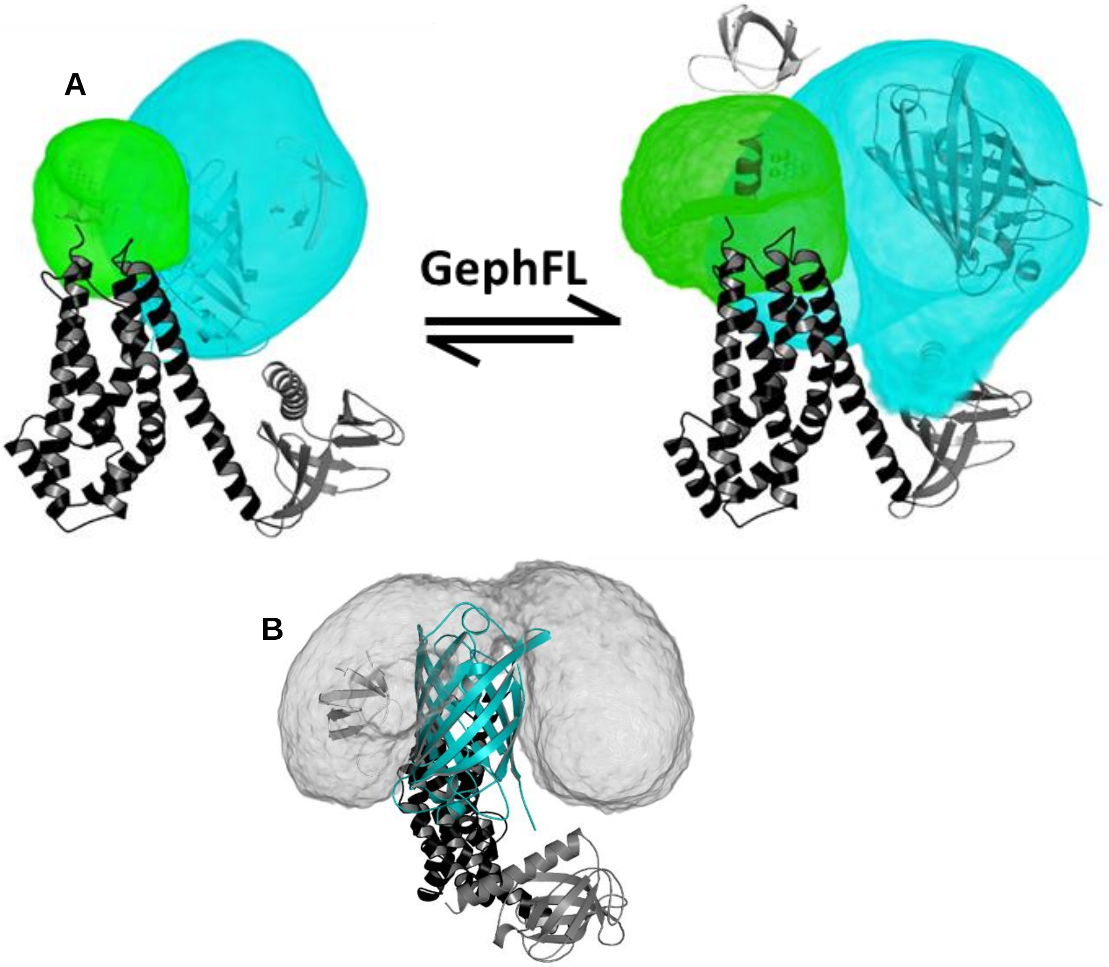


**Figure S12. Results of the FRET-restrained modelling.** (**A**) Closed state of CB, modeled based on PDB ID 4mt7 and the high FRET distances (left), and the open state of CB, modeled based on PDB ID 4mt7 and the low FRET distances. The DH domain is shown in black, PH in dark grey and SH3 in light grey. The cyan cloud depicts the accessible volume of CFP, whereas the green color represents the accessible volume of F99. (**B**) Accessible volume of the SH3 domain (grey density) in the open state conformation of CB. In the open state the intramolecular interaction of the SH3 and DH domains decreases and thus the mobility of the SH3 domain increases, which, in turn, brings the N-terminal FlAsH much closer to the C-terminally attached CFP leading to an increase in FRET efficiency.

**Table S1. Amino acid sequence of CB FRET sensors.** In each CB sensor, the residues highlighted in green and cyan represent the tetra-cysteine motif and CFP tag, respectively. The number in each name represents the position of the tetra-cysteine insertion in the CB protein. CFP is always at the C-terminus.

**F1$_{DA}$**

MCCPGCCQWIRGGSGMLITGDSIVSAEAVWDHVTELAFKAGDVIKVLDASNKDWWWGQIDDEEGWFPASFVRLWVNQ
EDGVEEGPSDVQNGHLDPNSDCLCLGRPLQNRDQMRANVINEIMSTERHYIKHLKDICEGYLKQCRKRRDMFSDEQLKV
IFGNIEDIYRFQMGFVRDLEKQYNNDDPHLSEIGPCFLEHQDGFWIYSEYCNNHLDACMELSKLMKDSRYQHFFEACRLL
QQMIDIAIDGFLLTPVQKICKYPLQLAELLKYTAQDHSDYRYVAAALAVMRNVTQQINERKRRLENIDKIAQWQASVLD
WEGDDILDRSSELIYTGEMAWIYQPYGRNQQRVFFLFDHQMVLCKKDLIRRDILYYKGRIDMDKYEVIDIEDGRDDDFN
VSMKNAFKLHNKETEEVHLFFAKKLEEKIRWLRAFREERKMVQEDEKIGFEISENQKRQAAMTVVSKGEELFTGVVPIL
VELDGDVNGHRFSVSGEGEGDATYGKLTLKFICTTGKLPVPWPTLVTTLTWGVQCFSRYPDHMKQHDFFKSAMPEGYV
QERTIFFKDDGNYKTRAEVKFEGDTLVNRIELKGIDFKEDGNILGHKLEYNYISHNVYITADKQKNGIKAHFKIRHNIEDG
SVQLADHYQQNTPIGDGPVLLPDNHYLSTQSALSKDPNEKRDHMVLLEFVTAAGITLGMDELYK

**F28$_{DA}$**

MQWIRGGSGMLITGDSIVSAEAVWDHVTCCPGCCELAFKAGDVIKVLDASNKDWWWGQIDDEEGWFPASFVRLWVNQ
EDGVEEGPSDVQNGHLDPNSDCLCLGRPLQNRDQMRANVINEIMSTERHYIKHLKDICEGYLKQCRKRRDMFSDEQLKV
IFGNIEDIYRFQMGFVRDLEKQYNNDDPHLSEIGPCFLEHQDGFWIYSEYCNNHLDACMELSKLMKDSRYQHFFEACRLL
QQMIDIAIDGFLLTPVQKICKYPLQLAELLKYTAQDHSDYRYVAAALAVMRNVTQQINERKRRLENIDKIAQWQASVLD
WEGDDILDRSSELIYTGEMAWIYQPYGRNQQRVFFLFDHQMVLCKKDLIRRDILYYKGRIDMDKYEVIDIEDGRDDDFN
VSMKNAFKLHNKETEEVHLFFAKKLEEKIRWLRAFREERKMVQEDEKIGFEISENQKRQAAMTVVSKGEELFTGVVPIL
VELDGDVNGHRFSVSGEGEGDATYGKLTLKFICTTGKLPVPWPTLVTTLTWGVQCFSRYPDHMKQHDFFKSAMPEGYV
QERTIFFKDDGNYKTRAEVKFEGDTLVNRIELKGIDFKEDGNILGHKLEYNYISHNVYITADKQKNGIKAHFKIRHNIEDG
SVQLADHYQQNTPIGDGPVLLPDNHYLSTQSALSKDPNEKRDHMVLLEFVTAAGITLGMDELYK

**F73$_{DA}$**

MQWIRGGSGMLITGDSIVSAEAVWDHVTMANRELAFKAGDVIKVLDASNKDWWWGQIDDEEGWFPASFVRLWVCCP
GCCNQEDGVEEGPSDVQNGHLDPNSDCLCLGRPLQNRDQMRANVINEIMSTERHYIKHLKDICEGYLKQCRKRRDMFS
DEQLKVIFGNIEDIYRFQMGFVRDLEKQYNNDDPHLSEIGPCFLEHQDGFWIYSEYCNNHLDACMELSKLMKDSRYQHF
FEACRLLQQMIDIAIDGFLLTPVQKICKYPLQLAELLKYTAQDHSDYRYVAAALAVMRNVTQQINERKRRLENIDKIAQW
QASVLDWEGDDILDRSSELIYTGEMAWIYQPYGRNQQRVFFLFDHQMVLCKKDLIRRDILYYKGRIDMDKYEVIDIEDG
RDDDFNVSMKNAFKLHNKETEEVHLFFAKKLEEKIRWLRAFREERKMVQEDEKIGFEISENQKRQAAMTVVSKGEELFT
GVVPILVELDGDVNGHRFSVSGEGEGDATYGKLTLKFICTTGKLPVPWPTLVTTLTWGVQCFSRYPDHMKQHDFFKSA
MPEGYVQERTIFFKDDGNYKTRAEVKFEGDTLVNRIELKGIDFKEDGNILGHKLEYNYISHNVYITADKQKNGIKAHFKI
RHNIEDGSVQLADHYQQNTPIGDGPVLLPDNHYLSTQSALSKDPNEKRDHMVLLEFVTAAGITLGMDELYK

**F99$_{DA}$**

MQWIRGGSGMLITGDSIVSAEAVWDHVTMANRELAFKAGDVIKVLDASNKDWWWGQIDDEEGWFPASFVRLWVNQE
DGVEEGPSDVQNGHLDPNSDCLCCPGCCQNRDQMRANVINEIMSTERHYIKHLKDICEGYLKQCRKRRDMFSDEQLKVI
FGNIEDIYRFQMGFVRDLEKQYNNDDPHLSEIGPCFLEHQDGFWIYSEYCNNHLDACMELSKLMKDSRYQHFFEACRLL
QQMIDIAIDGFLLTPVQKICKYPLQLAELLKYTAQDHSDYRYVAAALAVMRNVTQQINERKRRLENIDKIAQWQASVLD
WEGDDILDRSSELIYTGEMAWIYQPYGRNQQRVFFLFDHQMVLCKKDLIRRDILYYKGRIDMDKYEVIDIEDGRDDDFN
VSMKNAFKLHNKETEEVHLFFAKKLEEKIRWLRAFREERKMVQEDEKIGFEISENQKRQAAMTVVSKGEELFTGVVPIL
VELDGDVNGHRFSVSGEGEGDATYGKLTLKFICTTGKLPVPWPTLVTTLTWGVQCFSRYPDHMKQHDFFKSAMPEGYV
QERTIFFKDDGNYKTRAEVKFEGDTLVNRIELKGIDFKEDGNILGHKLEYNYISHNVYITADKQKNGIKAHFKIRHNIEDG
SVQLADHYQQNTPIGDGPVLLPDNHYLSTQSALSKDPNEKRDHMVLLEFVTAAGITLGMDELYK

**Table S2. Primers used in this study.**

| Primers | Forward (5'-3') | Reverse (5'-3') |
|---|---|---|
| CBSH3+ insertion | CTGGAAGTTCTGTTCCAGGGGCCCATGCAGTGGATTAGAG | TGGTGGTGGTGGTGCTCGAGTTACACAGTCATTGCGGCCTGTCTCT |
| CFP insertion | CAGGCCGCAATGACTGTGGTGAGCAAGGGCGAGGAG | TGGTGGTGGTGGTGCTCGAGTTACTTGTACAGCTCGTCCATG |
| tcM1 insertion | CCAGGGGCCCATGTGTTGCCCGGGCTGCTGTCAGTGGATTAGAGGCG | GTCGCCAGCCTTAAATGCCAACTCCCGGTTGGCCATGGTGACGTGATCCCATA |
| tCM28 insertion | AGTATGGGATCACGTCACCTGTTGCCCGGGCTGCTGTGAGTTGGCATTTAAGGCTGGC | GCCTCACAAAGCTGGCAGGAAACCATCCCTCCTCATCG |
| tCM73 insertion | AGCTTTGTGAGGCTCTGGGTGTGTTGCCCGGGCTGCTGTAACCAGGAGGATGGGGTGGAG | CTCCACCCCATCCTCCTGGTTACAGCAGCCCGGGCAACACACCCAGAGCCTCACAAAGCT |
| tCM99 insertion | ACTCAGACTGCCTCTGTTGCCCGGGCTGCTGTCAGAACCGGGACCAGAT | ATCTGGTCCCGGTTCTGACAGCAGCCCGGGCAACAGAGGCAGTCTGAGT |
| W24A mutation | GCTGAGGCAGTATGGGATCACGTCACC | GGTGACGTGATCCCATACTGCCTCAGC |
| E262A mutation | CCTTACAATTGGCCGAGCTCCTAAAGTA | TACTTTAGGAGCTCGGCCAATTGTAAGG |
| SH3 domain subcloning | TCTGGAAGTTCTGTTCCAGGGGCCCATGCTGATCACTGGAGATTCCAT | TGAACCAGGAGGATGGGGTGTAACTCGAGCACCACCACCACCACCA |
| CFP subcloning | GAAGTTCTGTTCCAGGGGCCCATGGTGAGCAAGGGCGAGGAGCTG | GGTGGTGGTGGTGGTGCTCGAGTTACTTGTACAGCTCGTCCATGCCG |

**Table S3:** Species-weighted average fluorescence lifetime ($\langle\tau\rangle$) and inter-dye distances ($R_i$) along with their relative species fractions ($x_i$) obtained from time-resolved FRET analysis for the F1 CB-FRET sensor in the absence ($F1_{D0}$), presence of FlAsH ($F1_{DA}$) alone and after incubation with $NL2_{icd}$, SH3 domain, full-length gephyrin (GephFL) and its domain variants. Species fractions are normalized such that $x_1 + x_2 + x_{noFRET} = 1$.

| Sample | $\langle\tau\rangle$ (±SD), [ns] | $E_{FRET}$ [%] | $R_1$ (±SD) [Å] | $x_1$ | $R_2$ (±SD) [Å] | $x_2$ | $x_{NoFRET}$ |
|---|---|---|---|---|---|---|---|
| $F1_{D0}$ | 3.10 (±0.04) | - | - | - | - | - | - |
| $F1_{DA}$ | 2.50 (±0.02) | 19 | 25.5 (±0.5) | 0.59 (±0.02) | 45.5 (±0.9) | 0.19 (±0.02) | 0.21 (±0.03) |
| $F1_{DA}$ +GephFL | 2.73 (±0.02) | 12 | 26.3 (±0.6) | 0.31 (±0.02) | 47.1 (±1.8) | 0.30 (±0.02) | 0.38 (±0.01) |
| $F1_{DA}$ +GephG | 2.51 (±0.03) | - | 26.8 (±0.6) | 0.43 (±0.01) | 44.5 (±0.6) | 0.29 (±0.01) | 0.27 (±0.01) |
| $F1_{DA}$ +GephLE | 2.67 (±0.01) | 14 | 26.3 (±0.47) | 0.37 (±0.01) | 49.6 (±0.47) | 0.25 (±0.01) | 0.37 (±0.03) |
| $F1_{DA}$ +GephE | 2.59 (±0.02) | 16 | 23.6 (±0.4) | 0.42 (±0.01) | 46.5 (±1.8) | 0.27 (±0.01) | 0.31 (±0.02) |
| $F1_{DA}$ + $GephE_{mm}$ | 2.85 (±0.01) | 8 | 24.5 (±0.2) | 0.26 (±0.02) | 43.7 (±0.3) | 0.16 (±0.02) | 0.57 (±0.01) |
| $F1_{DA}$ + $cytNL_{icd}$ | 2.60 (±0.02) | 16 | 25.7 (±0.12) | 0.44 (±0.01) | 46 (±0.62) | 0.23 (±0.01) | 0.32 (±0.01) |
| $F1_{DA}$ +SH3 | 2.66 (±0.02) | 14 | 26.9 (±0.88) | 0.42 (±0.03) | 49.3 (±0.9) | 0.27 (±0.04) | 0.29 (± 0.03) |

**Table S4:** Average fluorescence lifetime ($\langle\tau\rangle$) for $F1_{D0}$ in the absence and presence of full-length Gephyrin (GephFL), its domain variants (GephG, GephLE, GephE) and the dimer-deficient monomeric E-domain mutant ($GephE_{mm}$). The table also depicts the observed lifetime in $F1_{D0}$ in the presence of SH3 and $NL2_{icd}$.

| Sample | $\langle\tau\rangle$ (±SD), [ns] |
|---|---|
| $F1_{D0}$ | 3.10 (±0.04) |
| $F1_{D0}$ + GephFL | 3.11 (±0.02) |
| $F1_{D0}$ + GephG | 3.12 (±0.01) |
| $F1_{D0}$+ GephLE | 3.12 (±0.03) |
| $F1_{D0}$ + GephE | 3.11 (±0.01) |
| $F1_{D0}$+$GephE_{mm}$ | 3.15 (±0.01) |
| $F1_{D0}$+SH3 | 3.10 (±0.01) |
| $F1_{D0}$+$NL2_{icd}$ | 3.12 (±0.02) |

**Table S5:** Average fluorescence lifetimes ($\langle\tau\rangle$) of single ($F1_{smD0}$) and double mutant ($F1_{dmD0}$) CB FRET sensors, their FlAsH labeled counterparts $F1_{smDA}$ and $F1_{dmDA}$ in the absence and presence of full-length gephyrin (GephFL).

| Sample | $\langle\tau\rangle$ (±SD) ns | $E_{FRET}$ (%) |
|---|---|---|
| $F1_{smD0}$ | 3.12 (±0.02) | - |
| $F1_{smD0}$ + GephFL | 3.12 (±0.03) | - |
| $F1sm_{DA}$ | 1.15 (±0.03) | 63 |
| $F1sm_{DA}$ +GephFL | 1.98 (±0.03) | 36 |
| $F1dm_{D0}$ | 3.13 (±0.03) | - |
| $F1dm_{D0}$ + GephFL | 3.11 (±0.01) | - |
| $F1_{dmDA}$ | 1.16 (±0.02) | 63 |
| $F1_{dmDA}$ +GephFL | 2.05 (±0.05) | 34 |

**Table S6:** Average fluorescence lifetimes ($\langle\tau\rangle$), inter-fluorophore distances ($R_i$) and their relative species fractions ($x_i$) estimated from time-resolved FRET analysis for different CB-FRET sensors having the FlAsH moiety at positions 1, 28, 73, or 99 of CB in the presence of 100 µM full length gephyrin (GephFL). Species fractions are normalized such that $x_1 + x_2 + x_{noFRET} = 1$.

| Sample | $\langle\tau\rangle$ (±SD), [ns] | $E_{FRET}$ (%) | $R_1$ (±SD) [Å] | $x_1$ | $R_2$ (±SD) [Å] | $x_2$ | $x_{NoFRET}$ |
|---|---|---|---|---|---|---|---|
| $F28_{DA}$ | 2.54 (±0.01) | 18 | 25.9 (±1.1) | 0.47 (±0.02) | 48.1 (±0.4) | 0.25 (±0.02) | 0.28 (±0.01) |
| $F28_{DA}$ +GephFL | 2.64 (±0.01) | 15 | 26.9 (±0.2) | 0.33 (±0.01) | 47.4 (±0.3) | 0.29 (±0.01) | 0.38 (±0.01) |
| $F73_{DA}$ | 2.07 (±0.01) | 33 | 23.1 (±0.4) | 0.86 (±0.01) | 45.5 (±0.2) | 0.07 (±0.01) | 0.06 (±0.01) |
| $F73_{DA}$ +GephFL | 2.40 (±0.01) | 22 | 23.8 (±0.5) | 0.64 (±0.01) | 42.2 (±1.7) | 0.15 (±0.01) | 0.20 (±0.01) |
| $F99_{DA}$ | 2.28 (±0.02) | 26 | 22.9 (±0.3) | 0.86 (±0.01) | 52.4 (±1.5) | 0.05 (±0.01) | 0.08 (±0.01) |
| $F99_{DA}$ +GephFL | 2.60 (±0.01) | 16 | 24.1 (±0.2) | 0.59 (±0.01) | 42.5 (±0.8) | 0.12 (±0.01) | 0.29 (±0.01) |

**Table S7. Connectivity restraints**. Components (Comp.) 1 and 2 denote the respective structural components / models to be connected. The length in amino acid residues is converted into Å assuming a worm-like chain polymer behaviour. The width of the distribution, used as uncertainty in FPS, was determined as the 1σ-interval (68% are under the curve).

| Comp. 1 | Residue | Atom ID | Comp. 2 | Residue | Atom ID | Length [aa] | Length [Å] | width [Å] |
|---|---|---|---|---|---|---|---|---|
| SH3 | Trp 72 | 442 | DH | Asn 106 | 445 (4mt6), 10 (4mt7) | 35 | 39.4 | 9.5 |
| PH | Lys 439 | 3291 (4mt6) 2840 (4mt7) | CFP | Lys 3 | 1 | --- | 18.4 | 3.6 |
| FlAsH-1 | Pro 6 | 106 | SH3 | Val 18 | 1 | 17 | 26.7 | 6.5 |
| SH3 | Trp 72 | 442 | FlAsH-99 | Pro 6 | 106 | 28 | 35.0 | 8.5 |
| FlAsH-99 | Pro 6 | 106 | DH | Asn 106 | 445 (4mt6), 10 (4mt7) | 7 | 15.6 | 4.0 |

**Table S8: Experimental FRET-based restraints.** Components (Comp.) 1 and 2 denote the respective structural components / models to be connected. The mean distances and uncertainties were determined as described in Supplementary Methods.

| Comp. 1 | Residue number | Atom ID | Comp. 2 | Residue number | Atom ID | $\langle R_{DA,closed}\rangle$ [Å] | $\delta R_{DA,closed}$ [Å] | $\langle R_{DA,open}\rangle$ [Å] | $\delta R_{DA,open}$ [Å] |
|---|---|---|---|---|---|---|---|---|---|
| FlAsH-1 | Pro 6 | 106 | CFP | Leu 64* | 490 | 22.3 | 2.2 | 43.8 | 1.3 |
| FlAsH-28 | Pro 6 | 106 | CFP | Leu 64* | 490 | 23.4 | 1.4 | 49.5 | 1.5 |
| FlAsH-73 | Pro 6 | 106 | CFP | Leu 64* | 490 | 23.4 | 1.6 | 40.1 | 1.6 |
| FlAsH-99 | Pro 6 | 106 | CFP | Leu 64* | 490 | 21.8 | 1.1 | 40.5 | 1.7 |
